# Predictive learning with local plasticity in excitatory-inhibitory networks

**DOI:** 10.64898/2026.08.31.748357

**Authors:** Henrique Reis Aguiar, Matthias H. Hennig

## Abstract

Predictive coding is a powerful normative framework for understanding cortical computation, but it is still an open question how biologically plausible networks with local plasticity support predictive inference and representation learning. In this work we show that a recurrent excitatory-inhibitory circuit with purely local plasticity can perform predictive inference without explicit error representations. We establish a direct analytic link that shows that learning in these circuits requires the weights to remain on a consistency manifold where recurrent inhibition matches the inhibition required by the predictive coding objective. Using a closed-form derivation of the consistency condition, we derive a plasticity rule that maintains it exactly under a Gaussian prior. Under a non-Gaussian prior, we find the rule supports learning sparse, factorized features, such as edge detectors from natural images. We show empirically that a BCM-like rule with an activity-dependent threshold approximates this well, while other Hebbian-like rules tend to learn less accurate solutions because they keep the weights too far from this manifold. With recurrent excitation the networks acquire spatiotemporal features like direction selectivity, and the ability to complete partially observed sequences. Time-continuous learning then leads to the development of low-dimensional attractor-like structures and noise-driven replay. Overall, these results link predictive coding to local circuit plasticity, show it does not require an explicit prediction error representation, and suggest a normative role for BCM-like plasticity in excitatory synapses.

## Introduction

Excitatory-inhibitory (E-I) networks are local, recurrently connected neural circuits that are observed ubiquitously throughout the brain [Wilson and Cowan, 1972, Isaacson and Scanziani, 2011, Sanzeni et al., 2020]. Through the interaction of recurrent excitation and inhibition, these circuits can generate a rich repertoire of dynamical regimes, including discrete attractors [Hopfield, 1982, Krotov and Hopfield, 2016], continuous attractors [Amari, 1977, Hopfield, 2015], ring attractors [Zhang, 1996], oscillations [Buzsaki and Draguhn, 2004], and sequential activity [Rajan et al., 2016]. Such dynamics have been proposed to support functions as diverse as working memory, sensory integration, evidence accumulation, spatial navigation, motor control and much more [Khona and Fiete, 2022, Vyas et al., 2020]. However, the fundamental principles that determine how E-I networks self-organize into these functional circuits remain poorly understood.

Here, we look at these circuits through the lens of predictive coding, which views each circuit as learning an internal probabilistic model of its inputs, whether these are sensory signals or the activity of other neural populations, and using this model to infer their latent causes, complete missing information, and anticipate future observations [Barlow, 1989, Ackley et al., 1985, Hinton et al., 1995, Friston, 2005]. Within this framework, recurrent neural dynamics perform inference by reconciling incoming data with the current model, while synaptic plasticity optimizes the model to better fit the statistical structure of incoming data [Olshausen and Field, 1997, Rao and Ballard, 1999, Bogacz, 2017]. Predictive coding therefore offers a normative account linking neural dynamics, perception, and learning. However, it is not yet clear how the diverse, local synaptic plasticity mechanisms observed experimentally can collectively optimize the probabilistic objectives posited by this theory within realistic E-I circuits, and therefore we address this problem here.

Classical predictive coding implementations have been shown to learn neural representations that are also observed experimentally, supporting the idea that biological networks implement a similar kind of learning. For instance, from static natural images, these models learn orientation-selective receptive fields also found in the primary visual cortex of mammals [Olshausen and Field, 1997, Bell and Sejnowski, 1997]. These *factorized* representations are very useful since they disentangle the underlying components of variation (i.e. *factors*), therefore allowing a network to represent new data (i.e. combinations of these factors) not seen before during training [Hinton, 1986, Bengio et al., 2013]. However, algorithmic implementations to optimize this objective mostly rest on mechanisms that have not been observed experimentally in the brain, such as symmetric feedback weights [Hinton et al., 1995, Olshausen and Field, 1997], error neurons [Rao and Ballard, 1999] and non-local updates. More recent implementations propose complex models of neurons where dendritic compartments can compute prediction errors locally [Urbanczik and Senn, 2014, Sacramento et al., 2018, Mikulasch et al., 2023].

A parallel line of research has focused on studying models that can learn factorized representations while remaining faithful to biology. Early work proposed that adaptive recurrent inhibition coupled with local plasticity can lead to the learning of factorized representations [Földiak, 1990]. This model, often called the Hebbian/anti-Hebbian network, was later shown to learn orientation selectivity from static natural images in both firing-rate models [Falconbridge et al., 2006] and spiking networks [Triesch, 2004, Savin et al., 2010, Zylberberg et al., 2011]. More recently, firing-rate models were derived from non-likelihood objectives, such as similarity matching [Pehlevan et al., 2017] and non-negative matrix factorization [Pehlevan and Chklovskii, 2014], paving a way to understanding these networks from first principles. However, an exact link between these more plausible models and the principles of predictive coding theory is still lacking.

Here we clarify this link by showing that Hebbian/anti-Hebbian networks can learn to implement a predictive coding objective with biologically plausible local plasticity rules. Key to this is the requirement that the excitatory feed-forward weights are updated such that they remain on a *consistency manifold* where the learned inhibition stays matched to the optimal inhibition obtained from the predictive coding objective. We propose a new local plasticity rule for the excitatory weights, termed the *predictive* rule, which keeps the weights on this consistency manifold throughout learning, leading to optimal learning. In non-linear networks we find that a Bienenstock-Cooper-Munro-like (BCM) rule [Bienenstock et al., 1982], shown before to learn better representations in networks with static inhibition [Krotov and Hopfield, 2019], also approximates the behaviour of the predictive rule. By contrast, we find that although optimal for adapting the inhibitory weights, classic Hebbian plasticity applied to the excitatory weights pushes networks off the consistency manifold. We then include recurrent excitation to model temporal dependencies in the latent state and observe the development of spatiotemporal selectivity from natural video data and the emergence of sequence completion abilities. Moving closer to biology, we implement a full excitatory-inhibitory network which applies BCM-like plasticity in a continuous fashion. This network develops attractor-like structures which not only keep activity within low-dimensional manifolds but also lead to the replay of learned sequences when the network is stimulated with noise. Together, these findings show that neural circuits equipped with learning rules derived from a predictive coding objective can reproduce central features of cortical computation.

## Results

### Predictive learning with biologically plausible mechanisms

Predictive coding theory posits that cortical activity is constantly inferring latent causes of its inputs, and perception constitutes inferring latent variables in a generative model [Rao and Ballard, 1999] (Fig. 1A). In its most simple form, sensory data **x** ∈ ℝ^*n*^ is generated from latent causes **y** ∈ ℝ^*m*^ via some *generative* distribution *p*(**x**|**y**) which can be parametrized to be a simple linear model *D* ∈ ℝ^*n×m*^ (also called a dictionary) with Gaussian noise *ε*, such that

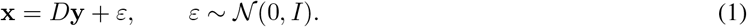

**Figure 1:**
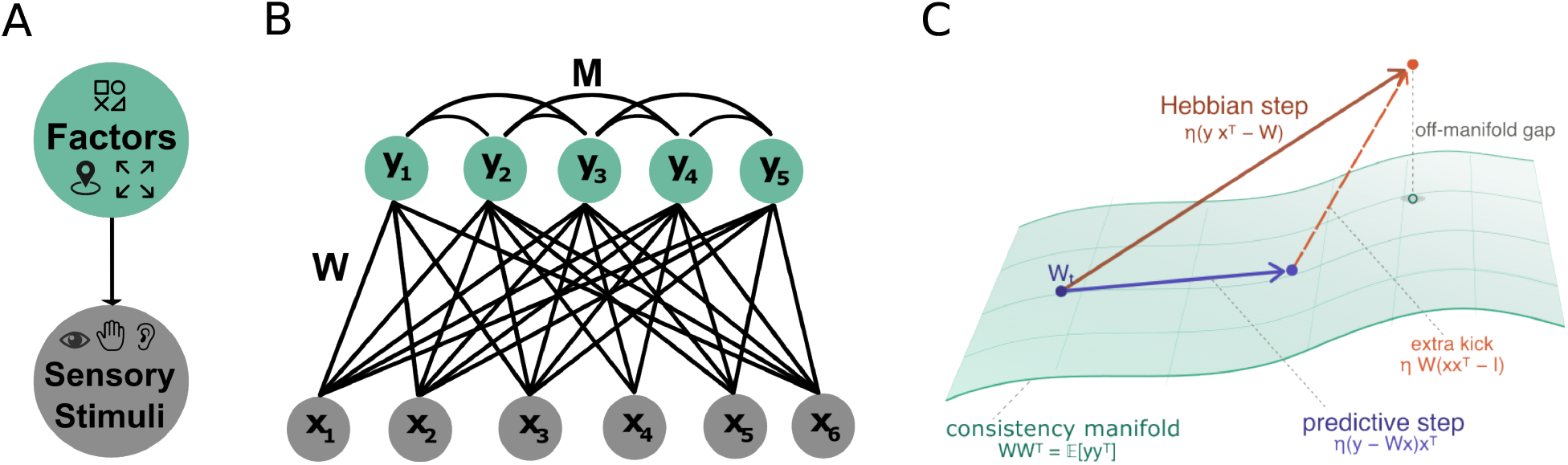
Predictive coding with lateral inhibition models. (A) Illustration of the generative modelling approach. (B) Illustration of the Hebbian/anti-Hebbian network with excitatory feed-forward connections *W* and inhibitory recurrent connections *M*. (C) Illustration of how a single weight update affects the trajectory of learning relative to the consistency manifold, which is defined as the non-linear subspace where *WW* ^*T*^ = E[**yy**^*T*^].

The latent code **y** is assumed to follow a *prior* distribution which is *factorized* into individual components *p*(*y*_*i*_) such that:

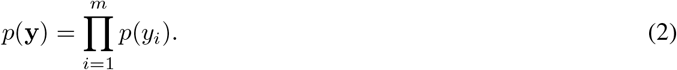

Assuming the individual prior distributions belong to an exponential family with energy *E*_*prior*_ (i.e. *p*(*y*_*i*_) ∝ exp(−*E*_*prior*_(*y*_*i*_))), one can write the full joint distribution as

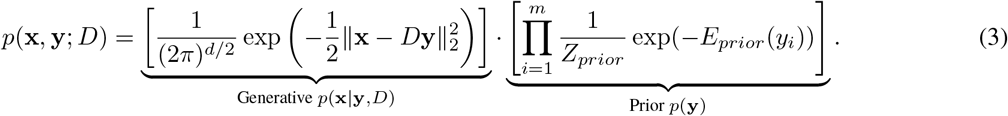

Standard Maximum Likelihood Estimation (MLE) seeks to learn *D* by maximizing the marginal likelihood *p*(**x**; *D*) = *p*(**x, y**; *D*) *d***y**. However, for some priors, this marginalization is analytically intractable. Predictive coding solves this by introducing an approximate posterior distribution *q*(**y**|**x**) which allows one to formulate a lower bound on the marginal log-likelihood, termed the evidence-lower bound (ELBO) [Jordan et al., 1998]. Further defining the approximate posterior to be a Dirac delta function, such that *q*(**y**) = *δ*(**y** − **y**^∗^), allows one to simplify the objective to that of minimizing the negative log of the joint probability or energy function [Olshausen and Field, 1997, Rao and Ballard, 1999], which for our model is defined as

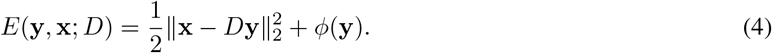

Here, we set the prior energy function to be 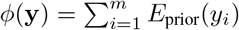 and removed the terms that do not include **y** or *D*. One can then optimize the joint energy with an expectation-maximization (EM) algorithm [Neal and Hinton, 1998, Olshausen and Field, 1997]. In the expectation step, the current parameter estimate *D*_*t*_ is fixed and the latent code **y**_*t*_ is inferred for the current sample **x**_*t*_. Performing maximum a posteriori (MAP) inference yields a point estimate for the expectation step, which is obtained by descending the gradient of *E* with respect to **y** until a stable state is found by

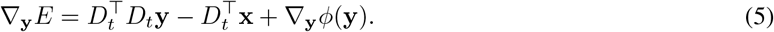

Note that this can be interpreted as the dynamics of a neural network with feed-forward inputs 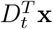 and lateral inhibitory connections 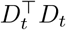. Once the latent code is obtained, the maximization step computes the next best estimate of the parameters *D*_*t*+1_ by taking a single descending step on the gradient of *E* with respect to *D*, such that

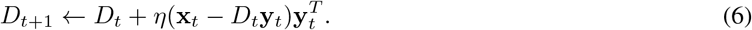

This is a well-known result from the literature. With a Laplace prior (which does not affect the weight update) it yields the locally competitive algorithm (LCA) introduced by [Rozell et al., 2008] as a biological implementation of the sparse coding objective [Olshausen and Field, 1997]. In this model, the weights for lateral inhibition and feed-forward connections are however tied, they do not change independently as in biological networks. Furthermore, the update rule for the dictionary *D* is implausible as it requires symmetric feedback connectivity to compute the reconstruction error. A more plausible approach is to define a feed-forward matrix *W* = *D*^⊤^ and a separate lateral inhibition matrix *M* ∈ ℝ^*m×m*^ which replaces *D*^⊤^*D*, leading to a circuit commonly known as the Hebbian/anti-Hebbian network (Fig. 1B). Previous work has either proposed these networks mechanistically [Földiak, 1990, Falconbridge et al., 2006], or derived them from an ICA objective [Savin et al., 2010] or other objective functions, such as similarity matching [Pehlevan et al., 2017] and non-negative matrix factorization [Pehlevan and Chklovskii, 2014].

For the dynamics of the Hebbian/anti-Hebbian circuit to be equivalent to the circuit that minimizes energy (Eqn. 5), one requires the outer product of the feed-forward weights to match the recurrent weights, such that *M* ≈ *WW*^⊤^. This could be enforced by adding an extra term to the energy that minimizes the Frobenius distance between these matrices (i.e. ||*M* − *WW*^⊤^||_*F*_). However, this term would yield biological implausible (non-local) rules for both *W* and *M* in the gradient computation for the maximization step. Another strategy is to set *WW*^⊤^ = E[**yy**^⊤^] which is true if the following *consistency conditions* are upheld: (1) the input is whitened (i.e. E[**xx**^⊤^] = *I*) and (2) the feed-forward weights predict the inhibitory dynamics such that *W* **x** = **y** for all **x** where **y** is the inferred latent code. If both (1) and are true, then we can write

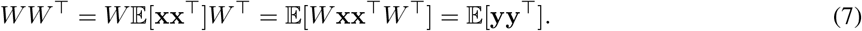

Substituting E[**yy**^⊤^] by *WW*^⊤^ in the original energy gives us an independent form for the lateral inhibition *M* which can be computed online (with an exponential decay *α*) via a simple Hebbian rule, such as

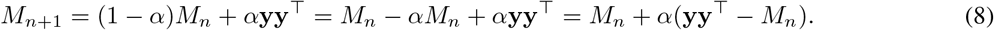

Substituting *D*^⊤^ by *W* in the original energy function (Eqn. 4), expanding it and replacing *WW*^⊤^ by our estimate of the second moment *M* = E[**yy**^⊤^] gives

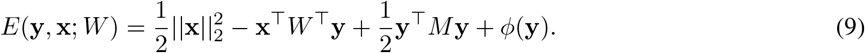

Note that this energy is equivalent to the original energy as long as the weights are inside the *consistency manifold* which is the manifold of the weight space where the consistency conditions are upheld (see Supplement for proof). Because the quadratic term is no longer dependent on *W*, performing gradient descent now becomes

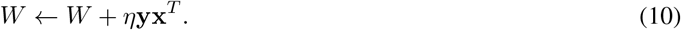

This weight update is however unstable since the expected increment is E[Δ*W*] = *η* E[**yx**^⊤^], leaving ∥*W*∥ to grow without bound. A penalty to prevent weight explosion can be added to the energy function in one of two ways:

a. **Weight normalization**. One may add a term to the energy function that penalizes large weights directly, such that

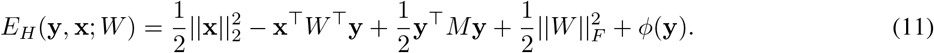 Taking the gradient of this energy with respect to the weights *W* will give us the classical *Hebbian* rule

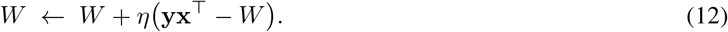
b. **Input normalization**. Alternatively, one can normalize the synaptic inputs, such that

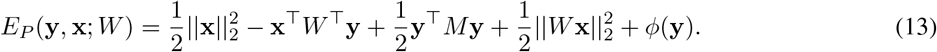

The gradient with respect to *W* gives us the *predictive* rule

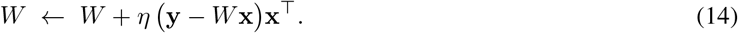

Under whitened inputs, E[*W* **xx**^⊤^] = *W*, so (12) and (14) have the same expected update E[Δ*W*] and the same fixed point (see derivations in the Supplement),

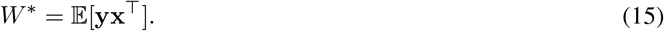

As long as the latent codes are inferred correctly, this shared fixed point inherits the same solution as the original objective in Eqn. 4 when the components have a constant norm (see proof 1.3 in Supplement). In order for the latent codes to be inferred correctly however, the consistency condition must hold for every step of the optimization procedure and not only at the fixed point. The predictive rule (14) is proportional to the consistency constraint error, defined as **e** = **y** − *W* **x**. Therefore, whenever the constraint is satisfied, the predictive update vanishes (i.e. **e** = 0), and whenever it is violated the update corrects it, leading the weights to always be inside (up to a small error for non-Gaussian priors) this consistency manifold (see proof 1.4 in Supplement). The Hebbian update (12) also applies the correction, but then pushes the weights away from the manifold by a factor of *W* (**xx**^⊤^ − *I*). This can be seen by rewriting the Hebbian gradient in terms of the predictive gradient

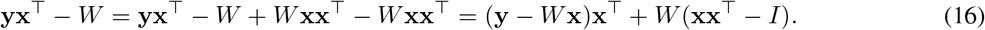

The second moment of the input (i.e. **xx**^⊤^) is only the identity on average. For any single sample, **xx**^⊤^ will be of rank one and therefore will kick the weights off the consistency manifold (see Fig 1C for illustration and proof 1.5 in Supplement). This is a problem for the Hebbian rule, because if the consistency conditions are not upheld at every update, the latent code is inferred under a perturbed energy, and the learning signal is computed from those perturbed codes, leading the learning trajectory to drift over time. It is worth noting that for a sparse prior with an over-complete representation, the consistency constraint can never be upheld exactly. However, as we find in experiments, the predictive rule keeps the weights close enough to the consistency manifold such that the network still finds optimal solutions.

### Predictive networks learn under both Gaussian and non-Gaussian priors

Setting the prior to be a standard Gaussian with zero mean and identity covariance yields a network with linear dynamics (Eqn. S10), which learns a version of the principal subspace of the input data [Tipping and Bishop, 1999, Pehlevan et al., 2017] (note the ordering is reversed, see Supplement S4). However, as shown above, this only works if the input data **x** is whitened, which leads the network to learn a trivial transformation since the input data is already following a Gaussian distribution with zero mean and identity covariance. Therefore, we test the network with non-whitened input data. As a first example, we draw a 2-dimensional input from a Gaussian distribution with zero mean and non-diagonal covariance (Fig. 2A, left column) and train a network with 2 units. Feed-forward weights are randomly initialized, and the lateral weights set to identity (Fig. 2A, first row). Learning only the lateral inhibition *M* with the Hebbian rule (with frozen feed-forward weights) leads the network to output a whitened distribution (i.e. the distribution defined by the prior above, see Fig. 2A, second row, right column). Subsequently, applying the predictive rule to learn the feed-forward weights leads the synaptic output **z** = *W* **x** to follow a whitened distribution (Fig. 2A, third row, middle column). Since the synaptic output is already following the distribution imposed by the prior, recurrent dynamics are not necessary any more, and inhibition reverts to the identity matrix.

**Figure 2:**
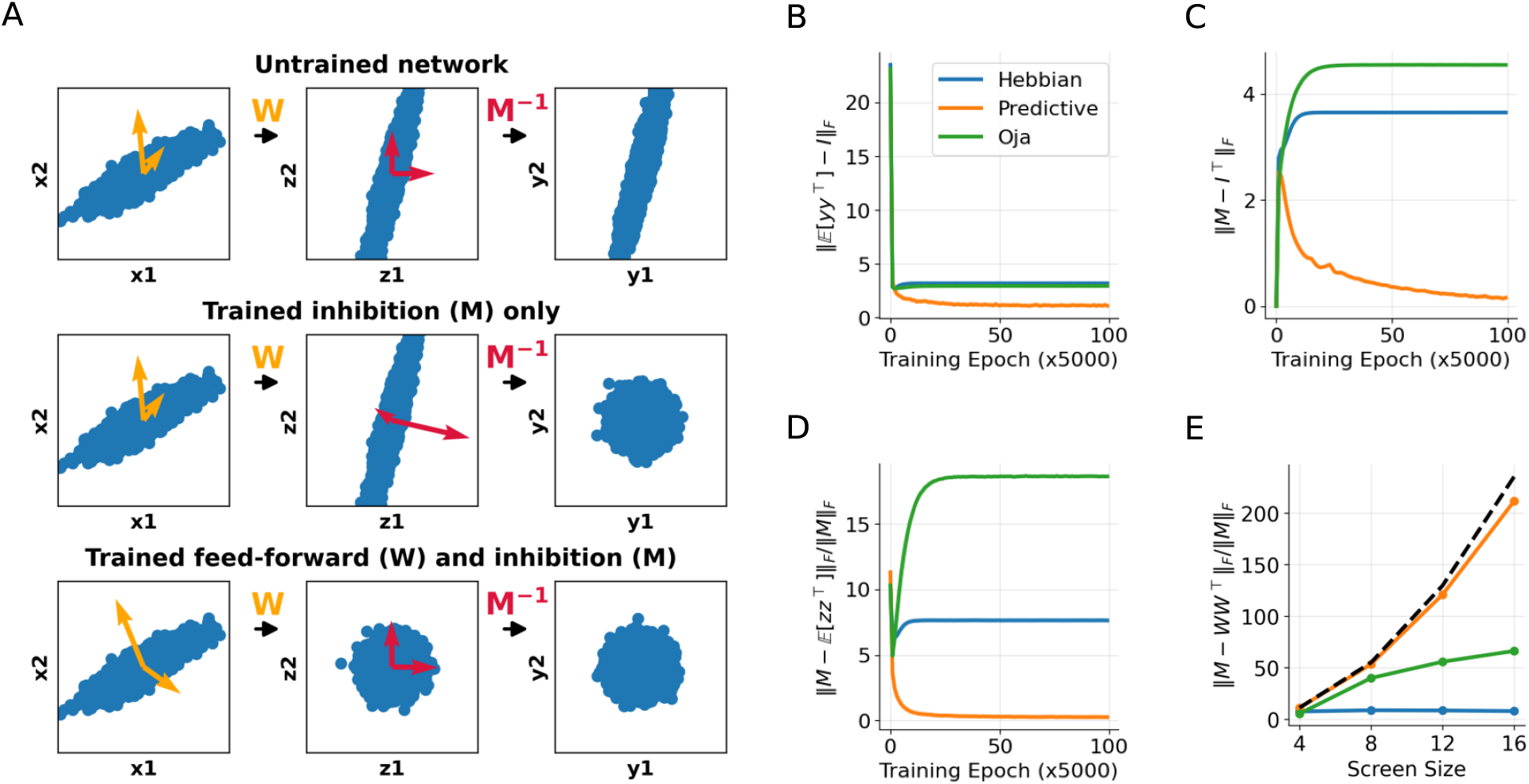
The predictive rule adapts to match inhibitory dynamics according to a predictive coding objective. **(A)** Example of transformations performed by a linear network with 2 input units (**x**_1_, **x**_2_) and two output units (**y**_1_, **y**_2_). The feed-forward weights *W* transform input into synaptic input **z** = *W* **x**, and the inverse inhibitory weights *M*^−1^ apply linear inhibitory dynamics to produce the network output **y** = *M*^−1^**z.**The yellow and red arrows illustrate the row vectors of each matrix, W and *M*^−1^) respectively, in the appropriate space. (B-D) A network of 16 units is simulated with the 4 by 4 non-whitened centered crossbar dataset. Feed-forward and recurrent weights are updated together throughout the simulation, with the inhibitory update rule always being Hebbian. (B) Distance between the encoded expected second moment E[**yy**^⊤^] and the identity matrix throughout the simulation. (C) Distance between the inhibition *M* and the identity matrix. (D) Distance between the inhibition *M* and the expected second moment E[**zz**^⊤^] = *W* E[**xx**^⊤^]*W*^⊤^ of the input. (E) Normalized consistency error (Eqn. S4) at the end of simulation with different screen sizes for different feed-forward plasticity rules. The number of neurons in each network equals the number of dimensions of the input (i.e. *m* = *n*, where *n* is the screen size squared). The black dashed line is the analytically derived error, emerging from the distance between E[**xx**^⊤^] and the identity matrix (see Supplement section S4 for details).

We validate these findings with the crossed-bars input of size 4 *×* 4 (see Supplement for details) and contrast the effect of the predictive rule (Eqn. 14) with other feed-forward local plasticity rules previously proposed: the Hebbian rule (Eqn. 12) [Pehlevan et al., 2017, Qin et al., 2023] and the Oja rule (Eqn. S22) [Oja, 1982, Pehlevan and Chklovskii, 2014]. We see that these rules learn a network that outputs a decorrelated response distribution (i.e. E[**yy**^⊤^] ≈ *I*), with the predictive rule slightly outperforming other plasticity rules (Fig. 2B). The biggest difference between the rules is the evolution of the inhibition (Fig. 2C), where most rules keep it far from the identity in order to keep activity decorrelated, while the predictive rule leads the feed-forward weights to learn this decorrelation transformation, making the synaptic input (i.e. **z** = *W* **x**) white and therefore leading the inhibition to drift back towards the identity since there is nothing else to decorrelate. For the predictive rule, the inhibition also satisfies the second part of the consistency condition (i.e. E[**yy**^⊤^] ≈ E[*W* **xx**^⊤^*W*^⊤^]), since the feed-forward weights can perfectly predict the stable state (Fig. 2D), while other rules do not achieve this result. However, non-white input does not allow the consistency constraint to be fully satisfied by the predictive rule (Fig. 2E). The consistency error is determined by the input spectrum, and the simulations agree well with its theoretical prediction (Fig. 2E, dashed line shows prediction; see Supplement S4 for derivation).

For a non-Gaussian prior, on the other hand, the input data is usually whitened before it is fed to the networks, since one wants the learning to focus on capturing higher order correlations [Olshausen and Field, 1997, Bell and Sejnowski, 1997]. Setting the prior to be exponential will yield a non-linear network where units are rectified in order to remain positive (Eqn. S15). This non-linearity, coupled with local plasticity, is often the recipe for finding sparse components of the input distribution [Földiak, 1990, Savin et al., 2010, Pehlevan and Chklovskii, 2014, Brito and Gerstner, 2016]. As above, we simulate networks where inhibition tracks the second moment of the stable state activity via Eqn. 8 and the feed-forward weights learn according to the different local plasticity rules introduced above. Here we additionally test a BCM-like rule [Bienenstock et al., 1982], which flips the sign of the update depending on whether post-synaptic activity is above or below the neuron’s mean activity (see Eqn. S27). Note that this rule is equivalent to the Hebbian rule in the Gaussian case because the mean activity of the neuron is zero. We also test the Földiák rule [Földiak, 1990] instead of the Oja rule, which is more appropriate for the non-Gaussian case where activity is strictly non-negative [Falconbridge et al., 2006]. Importantly, the input is strictly positive, which leads us to examine and compare networks that center the input (and therefore better fulfil the consistency constraint) versus ones that do not. Furthermore, we constrain the weight space at each update by setting the norm of the weight vector of each post-synaptic neuron to be 0.1 and also by rectifying the weights to remain non-negative, therefore moving the model closer to biology.

We simulated networks without centering under the crossed-bars stimulus (CROSS) dataset [Földiak, 1990] and observe that all rules learn the correct separation of the crosses into “stripes” and “bars” (Fig. 3A, top left). However, the final reconstruction error (Fig. 3D) as well as the number of active neurons differ between the learning rules. Activity histograms (Fig. 3A, top right) show that in networks where the feed-forward rule is a simple product of pre and post-synaptic activity (Hebbian and Földiák), all neurons remain active throughout the simulation. In contrast, the number of active neurons under BCM and the predictive rule is exactly equal to the number of factors required to encode the data (red vertical line in the histograms), which indicates that these rules are optimally reducing the redundancy of the encoding, which directly translates in improved reconstruction performance when compared to other plasticity rules (Fig. 3D). Although the consistency constraint cannot be fully satisfied because of lack of whitening and the weight overlap introduced by the non-Gaussian prior (which requires recurrence to explain away the overlapping features), we observe that all local plasticity rules tend to reduce the distance between the learned inhibition and the optimal inhibition during training (Fig. 3B). However, the predictive and BCM-like rules reduce the consistency error more than other rules (Figs. 3B,C) which leads to lower reconstruction errors (Fig. 3D). Centering the input yields lower the consistency errors for all rules when trained on the crossed-bar dataset (Supplement Fig. S4A). However, reconstruction for the predictive and BCM are lower for non-centered networks (Supplement Fig. S4B).

**Figure 3:**
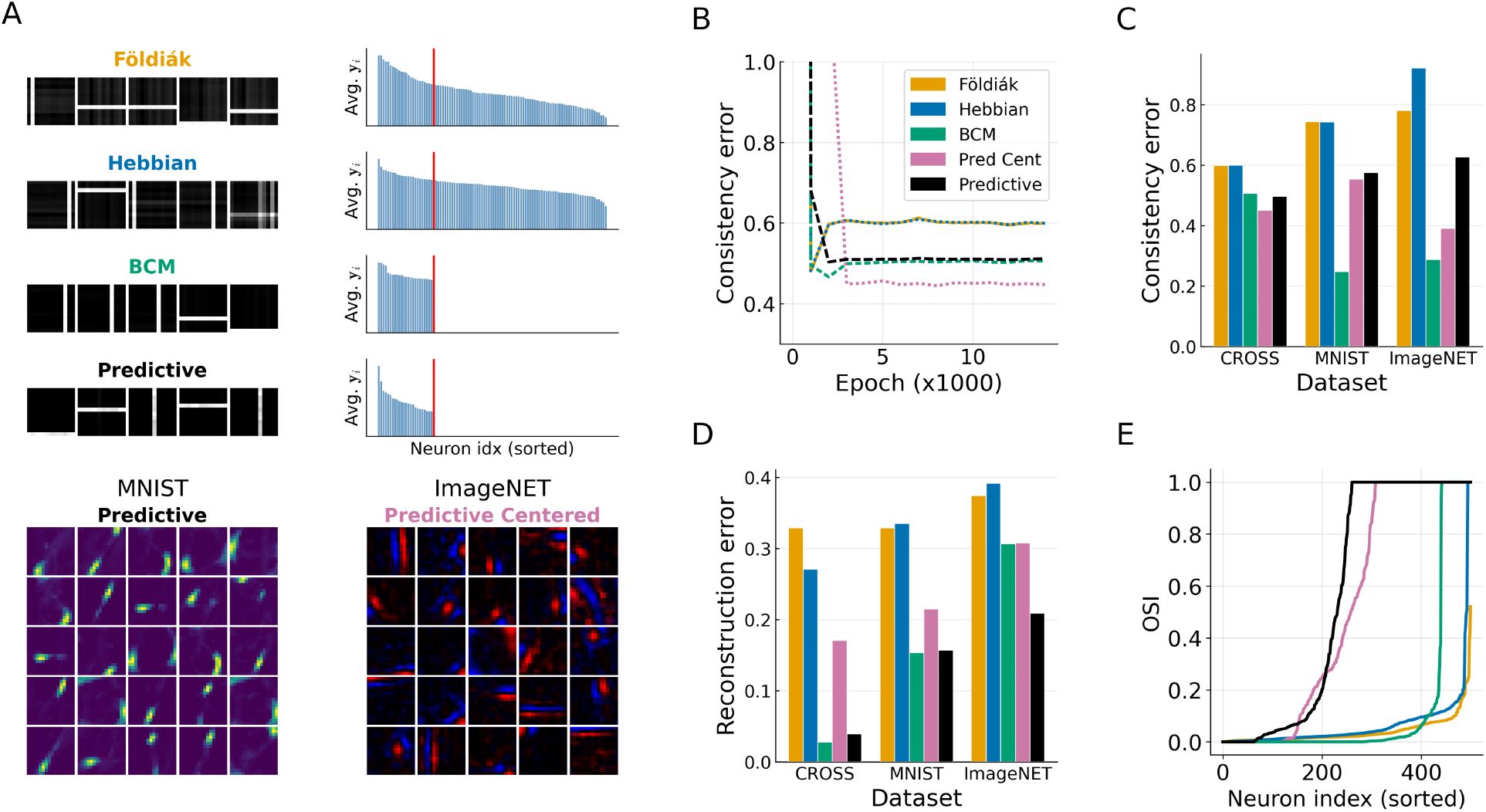
Consistent networks learn better solutions under non-Gaussian priors. (A) Receptive fields and activity histograms for networks trained under different feed-forward learning rules. Red lines in activity distributions mark the number of factors in the input distribution. Top 4 networks are learned under the crossbar dataset without weight normalization. Bottom left network is trained on the MNIST dataset and bottom right is trained on the ImageNET on-off patches dataset where both the input and weights are normalized at every update. (B) Normalized consistency error (Eqn. S4) for different feed-forward plasticity rules throughout training on the crossbar dataset. (C) Normalized consistency error at the end of the simulation for different training datasets and feed-forward plasticity rules. Experiments were run with 5 different seeds, the mean is plotted and the variability is on average 4.9 *×* 10^−7^ across rules. (D) Reconstruction error between the input **x** and the reconstruction from the stable state 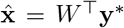. Experiments were run with 5 different seeds, the mean is plotted and the variability is on average 4.7 *×* 10^−5^ across rules. (E) Orientation selectivity index (OSI) for networks trained on ImageNET patches with different feed-forward rules.

We validate these findings the MNIST and ImageNET datasets. For MNIST we used cropped 16 *×* 16 patches, while for ImageNET, random patches were cropped, whitened and split into on and off channels (see Supplement S6). We observe the same pattern across these datasets (Fig. 3C-D), suggesting that both BCM and predictive rules can maintain the weights close enough to the consistency manifold as to find good solutions that minimize the reconstruction error and learn appropriate receptive fields (see Fig. 3A bottom and Supplement Figs. S5 and S6). Finally, we observe that the predictive rule learns broader orientation selectivity under both centered and non-centered inputs, compared to the other rules (Fig. 3E, see Supplement Fig. S6 for examples).

### Delayed recurrent excitation induces sequential learning

In addition to recurrent inhibition, the cortex has extensive recurrent excitation, which is suggested as the mechanism underlying the generation of sequential neural activity [Hopfield, 2015, Douglas et al., 1995]. To include recurrent excitation in our model, we propose that the current hidden state **y**_*t*_ has generated both the current input **x**_*t*_ ∈ ℝ^*n*^ and the previous hidden state **y**_*t*−1_ ∈ ℝ^*m*^. As before we assume that *p*(**y**_*t*_) follows a factorized non-Gaussian distribution. Then the generative process can then be formulated by concatenating the input and previous hidden state, such that [**x**_*t*_, **y**_*t*−1_] ∈ ℝ^*n*+*m*^, therefore obtaining

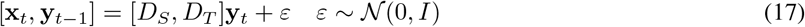

Here, the model for the spatial factors *D*_*S*_ ∈ ℝ^*n×m*^ and the model for the temporal factors *D*_*T*_ ∈ ℝ^*m×m*^ are also concatenated, such that [*D*_*S*_, *D*_*T*_] ∈ ℝ^(*n*+*m*)*×m*^, and therefore can be learned by minimizing the energy function

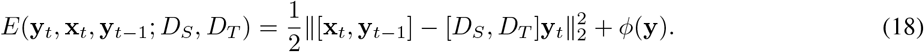

We assume the weights are kept within a consistency manifold as before, such that [*D*_*S*_, *D*_*T*_]^⊤^[*D*_*S*_, *D*_*T*_] = E[**yy**^⊤^]. We set the prior to be exponential, 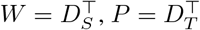 and *M* = E[**yy**^⊤^], and obtain the network dynamics,

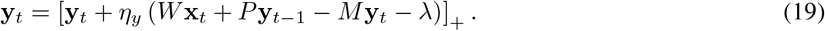

Note that this dynamics can be interpreted as having slow excitation, because one first needs to obtain the stable state for the current hidden state by keeping the inputs **x**_*t*_ and **y**_*t*−1_ fixed, where **y**_*t*−1_ is the previous hidden stable state (illustrated in Fig. 4A). The inhibitory plasticity rule is the online estimate of E[**yy**^⊤^] with an exponential decay (Eqn.8). As the model is a concatenation of the feed-forward *W* and recurrent *P*, we can update them individually using the predictive rule proposed above (Eqn. 14), such that

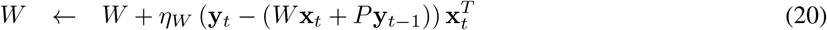

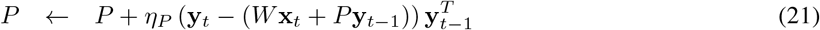

**Figure 4:**
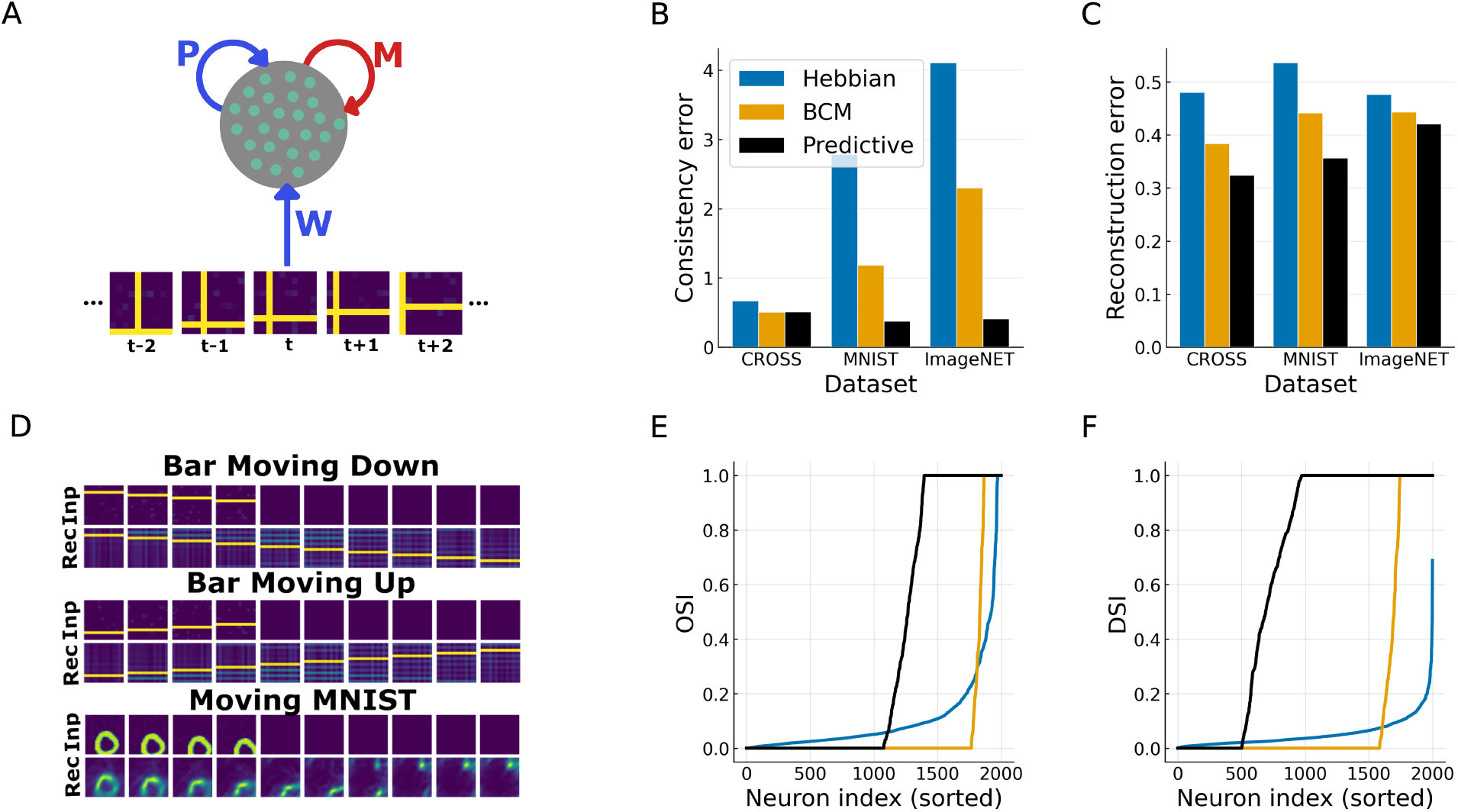
A sequence model learns spatio-temporal factors. (A) The sequence model adds slow recurrent excitation *P* to the Hebbian/anti-Hebbian network. (B) Normalized consistency error (Eqn. S4) for the sequential model at the end of the simulation for different excitatory plasticity rules and varied sequential datasets. (C) Sequence completion error for different excitatory plasticity rules. (D) Examples of sequence completion. Inputs were blanked from the fifth time step onwards, but decoded activity shows continuation of the previous movement pattern. (E) Orientation selectivity index (OSI) for networks trained on the CATCAM dataset with different feed-forward plasticity rules. (F) Direction selectivity index (DSI) for networks trained on the CATCAM dataset with different feed-forward plasticity rules.

We test our model on the moving crossed-bars, where the two bars move in independent directions (see Fig. 4A and Supplement Fig. S2). After learning, we observe the model is able to complete elementary sequences that continue the input sequence after input blanking (Fig. 4D). This shows the model has disentangled the spatio-temporal dynamics into independent, direction-selective factors. The sequence completion ability, measured as the error between the original and reconstructed sequence, emerges for both predictive and BCM rules (Fig. 4C). As before, the consistency error follows the same pattern across learning rules (Fig. 4B).

We validate these findings on the moving MNIST dataset and the CATCAM dataset which mimics animal visual experience [Betsch et al., 2004] (see Supplement for details). We first observe sequence completion emerging on the moving MNIST dataset trained with the predictive rule (Fig. 4D). Across datasets, we also observe the same pattern of consistency and reconstruction which shows that both predictive and BCM rules learn better solutions when applied to the excitatory weights (see Fig. 4B and 4C). We further analyse the orientation and direction selectivity of networks trained under the CATCAM dataset, and observe that both predictive and BCM rules learn highly selective cells, while selectivity for the Hebbian rule remains relatively low for all neurons, particularly the direction selectivity (Fig. 4E,F).

### Continuous networks develop attractor-like structure

The models studied above all had a caveat, they were temporally ‘discrete’: they receive one sample at a time, allow the dynamics to settle, and then apply the plasticity update. This is the basic principle of the expectation-maximization algorithm, but biological networks do not implement this synchronized flow of computation. Instead, stimuli enters as a continuous data stream, and plasticity events happen asynchronously, in pair with the activity dynamics. To address this gap, we propose a continuous excitatory-inhibitory model which learns from streams of data and applies synaptic updates in pair with the activity dynamics (see Methods and Supplement for details). These updates are gated by post-synaptic activity, following the work in Aguiar and Hennig [2024], which prevents post-synaptic neurons with recent plasticity updates or with low-firing rate from updating their incoming weights. In this continuous model we also introduce a separate population of inhibitory neurons in order to satisfy Dale’s principle [Dale, 1935] (Fig. 5A). Furthermore, following our theory, we use Hebbian plasticity for the inhibitory recurrent circuit and the BCM rule for the excitatory weights which, as shown above, performs a similar function to the predictive rule in terms of keeping the weights close to the consistency manifold.

**Figure 5:**
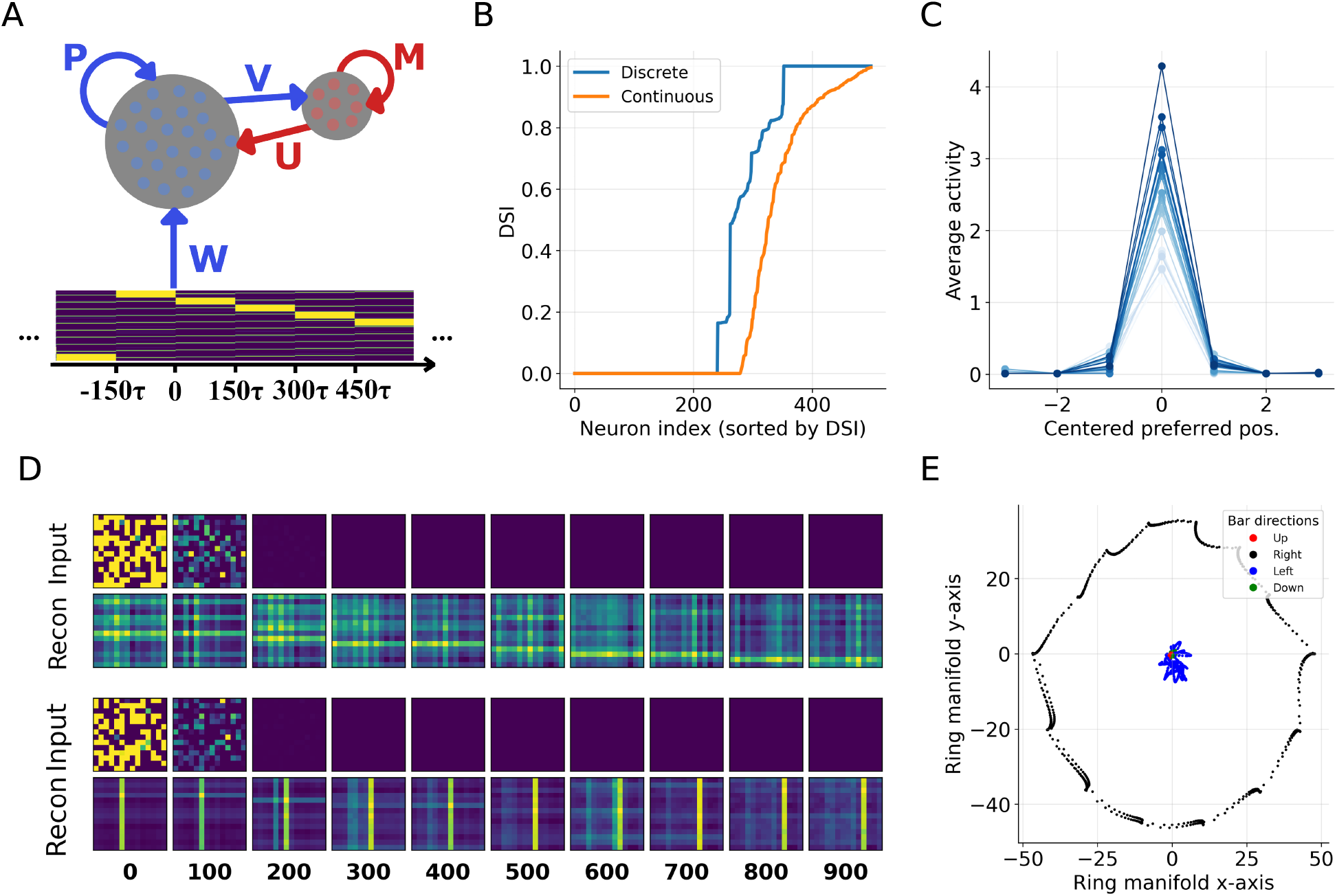
Continuous networks develop attractor-like properties when trained with sequential stimuli. (A) Illustration of the continuous E-I model with separate populations of excitatory (**y**) and inhibitory (**z**) neurons. The model is fed a flattened moving cross, where each frame is held for 150*τ* where *τ* is the time constant. (B) Direction selectivity index across the four possible directions of the discrete and continuous models when both are trained on the moving cross dataset with the BCM plasticity rule. (C) Tuning curves for different neurons in the continuous model, centered at the bar position which produces the highest response for each neuron. (D) Examples of spontaneous replay when the continuous model is briefly stimulated with random noise (inputs shown in the top rows). (E) Continuous model activity projected to a low-dimensional manifold when a bar moving in a specific direction is presented. We obtain one sequence of size 500 for each direction and plot the linearly projected activity for the whole sequence. Different colours represent activity points obtained for different sequences.

We simulate this model with a video stream of moving crosses that change uniformly every 150 simulation steps (Fig. 5A). We observe that, similarly to the discrete model, the continuous model develops direction selective cells (Fig. 5B) and learns to perform sequence completion (Supplement Fig. S3). Another interesting property emerging from the continuous model is that the spatial factors (i.e. bar position) are encoded by multiple neurons in a smooth fashion, resembling observed neural tuning curves rather than detectors for individual factors (Fig. 5C).

Stimulating the trained model with a burst of random noise leads to the replay of the spatio-temporal factors learned during training (Fig. 5D). The examples in Fig. 5D illustrate that the network starts by representing multiple elements of the same sequence (i.e. parallel bars), but as the sequence evolves, bars are removed through inhibition, leaving only a single element of the sequence at a time. This suggests that recurrent excitation and inhibition has developed to follow a ring attractor structure, which restricts activity to represent a single value of a variable at a time. To show this, we construct a linear transformation (see Supplement for details) which exposes one of these sequences (bar moving right) in 2D activity space. A projection of network activity obtained from playing different sequences shows a ring-like structure for this sequence, while activity from other sequences collapses around the origin, indicating that this activity inhabits other orthogonal subspaces (Fig. 5E).

## Discussion

In this work we show analytically that a recurrent inhibitory network with biologically plausible local plasticity can implement an EM-like procedure for learning a linear latent-variable model. Informally, this works if the feed-forward weights successfully predict the action of the recurrent inhibition between the encoding units, and when plasticity between the encoding units whitens their activity. Importantly, the network can only infer correct latent codes if the feed-forward weights are constrained to remain on a *consistency manifold*, a subset of the weight space where the independently learned inhibition is matched to the inhibition that originates from the predictive coding objective. Zylberberg et al. [2011] addressed this through through a Lagrange multiplier enforcing de-correlation through inhibition. By contrast, here we derived a predictive feed-forward plasticity rule that, for the linear Gaussian case, exactly maintains this condition because the update vanishes when the consistency condition holds and corrects deviations from the manifold. This rule is formally identical to the perceptron delta rule, but applied to the recurrently inferred latent code instead of an externally supplied label.

The network dynamics update allows specifying different priors for the encoding units. Consistent with previous work [Oja, 1982], we showed that a Gaussian prior yields linear dynamics and recovers a (reverse) principal subspace of non-whitened inputs, while an exponential prior recovers sparse, factorized directions like those found by normative models such as sparse coding and ICA [Olshausen and Field, 1997, Bell and Sejnowski, 1997, Hyvärinen and Oja, 2000, Falconbridge et al., 2006]. Under a non-Gaussian prior, however, the consistency condition can no longer be satisfied exactly. Nevertheless, we find empirically that the predictive rule and an experimentally supported BCM-like rule [Bienenstock et al., 1982, Kirkwood et al., 1996, Lim et al., 2015] both provide good approximations while the Hebbian and Földiák rules achieve poorer reconstructions. Furthermore, we show that the consistency condition can be approximately maintained when the weights are constrained to be non-negative, making the model compatible with Dale’s principle. It is interesting to note that the whitening of the code can also be achieved through gain modulation [Duong et al., 2023]. This is a biologically plausible mechanism for fast adaptation, while the factorisation can still be learned via synaptic plasticity. This could therefore be added to the continuous model to achieve the same outcome without need to re-learn inhibitory weights following distribution shifts.

Our framework offers an alternative to predictive coding models with explicit representation of mismatches between sensory activity and top-down predictions. These include error neurons [Rao and Ballard, 1999, Friston, 2005, Bastos et al., 2012], or errors represented within separate anatomical compartments of pyramidal cells [Sacramento et al., 2018, Mikulasch et al., 2021] or in the membrane potential in spiking networks [Brendel et al., 2020]. Although such models reproduce experimental observations including end stopping, repetition suppression, and mismatch responses [Rao and Ballard, 1999, Walsh et al., 2020, Mikulasch et al., 2023], it is still unclear whether the predicted explicit error representations exist, or if they arise from recurrent network dynamics. By contrast, in this work, prediction errors are absorbed in the local excitatory plasticity rule as the discrepancy between recurrently inferred activity and its feed-forward prediction. This replaces explicit error representation with a constraint on synaptic plasticity.

We extended the model to temporal inputs, yielding an excitatory-inhibitory circuit that exhibits spatio-temporal factorisation, such as the emergence of direction selectivity for natural video, and sequence completion. However, like previous temporal models [Shumway and Stoffer, 1982, Millidge et al., 2024], the discrete implementation alternates between inference and plasticity. This is difficult to reconcile with ongoing neural activity and can limit learning of processes evolving at multiple time scales.

We therefore proposed a continuous, sequential version of the model which implements an asynchronous update mechanism previously proposed by Aguiar and Hennig [2024], where plasticity and recurrent excitation evolve in pair with the inhibitory dynamics. A key assumption in this model, use-dependent refractoriness of plasticity, has experimental support [Kramár et al., 2012, Flores et al., 2025]. This model developed more realistic smooth tuning curves instead of the tuning to single factors in the discrete model, and spontaneous, noise-induced replay of previously seen sequences. This mirrors replay in the hippocampus [Wilson and McNaughton, 1994, Davidson et al., 2009], and spontaneous and induced spatio-temporal activity patterns in the visual cortex [Kenet et al., 2003, Carrillo-Reid et al., 2015, Gavornik and Bear, 2014, Xu et al., 2012]. We also observe population activity following low-dimensional trajectories, with different sequences near-orthogonal in neural space, consistent with low-dimensional attractor dynamics widely observed in neural populations [Churchland et al., 2012].

A limitation of the model is that the inference dynamics represent a single estimate of the latent state rather than posterior uncertainty or alternative latent explanations. As a result, plasticity has to rely on the inferred maximum a posteriori (MAP) code, which may not work well for ambiguous or partially observed inputs. Due to the point-estimate approximation, inhibition only tracks MAP statistics, which may lead to poor convergence. Anecdotally, we observed such behaviour in the form of winner-take-all dynamics and silent units under some parameter settings, while representation learning in the brain is reliable and robust. Moreover, an exact equivalence to the predictive-coding energy requires centered, whitened inputs and maintenance of the consistency condition throughout learning, which is only approximate for non-Gaussian priors, non-negative weights, over-complete representations and continuous dynamics. In the continuous model the same restrictions apply, but the latent trajectories may be interpreted as an online filtering estimate. This may remove the need for complete relaxation of the inference dynamics, and temporal continuity might instead stabilize changes between latent states, which are abrupt and less predictable in the discrete model.

Taken together, these results suggest that the computational role of local plasticity in excitatory-inhibitory networks is not only to decorrelate or sparsify activity, but to also hold the circuit in a regime where its own recurrent dynamics implement inference under an internal generative model. Then prediction errors do not have to be explicitly encoded, but can be expressed in the synaptic update. Consistent with this principle, simulations of spiking recurrent networks have shown that properly balanced inhibition is essential for the development of feature selectivity [Sadeh et al., 2015], and that inhibitory plasticity can act homeostatically to maintain sparse firing distributions [Schulz et al., 2021]. A central experimental prediction is then that excitatory plasticity should depend on the mismatch between recurrently inferred and feed-forward predicted activity. Consistent with this idea, slice experiments showed that inhibitory and excitatory plasticity can be inversely related in the same neuron [Wang and Maffei, 2014]. This may be testable in sensory systems, where manipulations of recurrent inhibition should alter the magnitude or sign of excitatory plasticity and should alter the selectivity and sparseness of learned representations.

## Methods

### Discrete model inference

To perform the latent code inference, network updates with inputs **x** ∈ ℝ^*n*^ and hidden units **y** ∈ ℝ^*m*^ were run as **y**_*t*+1_ = **y**_*t*_ +*τ*(*W* **x**−*M* **y**_*t*_ −**y**_*t*_) for the Gaussian model and **y**_*t*+1_ = [**y**_*t*_ +*τ*(W **x**−*M* **y**_*t*_ −*λ***1**)]+ for the exponential model with rate *λ. W* are feed-forward excitatory weights, *M* are recurrent inhibitory weights, *τ* = 0.01 is the integration step size, [·]_+_ enforces the non-negativity of the exponential prior and dynamics are iterated for 1000 steps. For the sequential model, we use an exponential prior.

### Plasticity rules

Following the inference step, a plasticity update is applied. The inhibitory weights *M* are always updated with the Hebbian rule (Eqn. 8). For the feed-forward weights *W* we compare the *predictive* rule (Eqn. 14), the Hebbian rule with weight decay (Eqn. 12), Oja’s rule (Eqn. S22), the Földiák rule (Eqn. S24), and a BCM rule with a first-order sliding threshold (Eqs. S26–S27). For the MNIST and ImageNET datasets, weights are constrained to be of norm 0.1, such that ||**w**_*i*_|| = 0.1, where **w**_*i*_ is the *i*th row of *W*.

### Continuous excitatory–inhibitory model

The continuous model has excitatory **y** ∈ ℝ^*m*^ and inhibitory **z** ∈ ℝ^*p*^ hidden neuron populations and runs plasticity in tandem with the activity dynamics on a continuous data stream. Inference dynamics in the excitatory population evolve according to **y**_*t*+1_ = [**y**_*t*_ + *τ* (**Wx**_*t*+1_ + **P***K*(**y**_*t*_, …, **y**_*t*−*β*_) − **Uy**_*t*_)]_+_ with feed-forward weights **W** ∈ ℝ^*m×n*^, recurrent excitatory weights **P** ∈ ℝ^*m×m*^ and temporal kernel *K*: ℝ^*mβ*^ → ℝ^*m*^ which integrates activity from the previous *β* ∈ N_+_ time steps. The inhibitory population evolves according to **z**_*t*+1_ = [**z**_*t*_ + *τ* (**Vy**_*t*+1_ − **Mz**_*t*_)]_+_, where *V* ∈ ℝ^*p×m*^ are the excitatory to inhibitory weights, *U* ∈ ℝ^*m×p*^ are the inhibitory to excitatory weights and *M* ∈ ℝ^*p×p*^ are the recurrent inhibitory weights, all updated according to the classic Hebbian rule (Eqn. S19). Both *W* and *P* are updated according to the BCM rule (Eqn. S26–S27) Plasticity updates are applied continuously with a stochastic threshold and refractory period following Aguiar and Hennig [2024].

### Stimuli

We use four stimulus families: (i) Crossed-bars on a 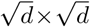 grid with each of the bars present independently or moving independently; (ii) a single moving bar (either horizontal or vertical) for testing OSI and DSI of different networks; (iii) MNIST digits [LeCun et al., 1998], either static in a 16 *×* 16 frame or moving in a 32 × 32 frame at a constant velocity; and (iv) for static natural images we use 16 *×* 16 ImageNet patches [Fei-Fei et al., 2009] whitened and split into two (on-off) channels, for natural video we crop a 16 *×* 16 patch from the CATCAM dataset [Betsch et al., 2004] whitened and split into two (on-off) channels. Unless stated, inputs are strictly positive and presented without centering. For the MNIST and ImageNET datasets, inputs are constrained to have a norm of 3, such that ||**x**_*i*_ || = 3 for all *i* in the dataset.

### Metrics

For static stimuli we report the mean cosine reconstruction error (i.e. 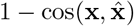 over 50 held-out inputs, with 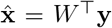. For sequential stimuli we measure extrapolation: the first half of each sequence is presented, the second half is replaced by blank input while the excitatory recurrent dynamics continue, and we report the per-frame cosine error with the original sequence 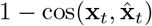 averaged over frames and 25 test sequences. Orientation (Eqn. S48) and direction (Eqn. S49) selectivity is measured with the synthetic stimuli for networks trained on naturalistic datasets. Throughout every simulation we also compute the distance of the weights from the consistency manifold, termed the consistency error (Eqn. S4).

## Acknowledgements

HRA was funded by an Engineering and Physical Sciences Research Council (EPSRC) DTA studentship. We thank Professor Chris Williams for valuable feedback on an early draft.

## Notes

### Competing Interest Statement

The authors have declared no competing interest.

