## Supplemental Material for "Predictive learning with local plasticity in excitatory-inhibitory networks"

### Supplemental notes

#### S1 Predictive coding optimization

**Data.** Inputs are vectors $\mathbf{x} \in \mathbb{R}^n$ drawn from a distribution where data is *centred* and *whitened*, such that $\mathbb{E}[\mathbf{x}] = 0$ and $\mathbb{E}[\mathbf{x}\mathbf{x}^\top] = I$.

**Model.** The matrix $W \in \mathbb{R}^{m \times n}$ stores $m$ learned feature vectors (*atoms*) as its rows $\mathbf{w}_1, \ldots, \mathbf{w}_m$, which we assume span $\mathbb{R}^n$, such that $m \geq n$. For each input, the *code* $\mathbf{y} \in \mathbb{R}^m$ is the minimizer of the substituted input normalized energy defined in equation 9 with respect to the code.

**Learning.** Let $\phi(\mathbf{y}) = \sum_i \phi_i(y_i)$ be the prior distribution, assumed to be continuous at the origin and normalized such that $\phi_i(0) = 0$. The reconstruction is $\hat{\mathbf{x}} = W^\top \mathbf{y}$, and the original predictive coding problem can be stated as follows:

$$E(\mathbf{y}, \mathbf{x}) = \tfrac{1}{2}\|\mathbf{x} - \hat{\mathbf{x}}\|^2 + \phi(\mathbf{y}), \qquad F(W) = \mathbb{E}\big[\min_{\mathbf{y}} E(\mathbf{y}, \mathbf{x})\big]. \tag{S1}$$

Solving this optimization problem via the expectation-maximization procedure yields a circuit with lateral inhibition and local plasticity. However, the lateral inhibition in this circuit is directly computed from the feed-forward weights, which is biologically implausible. In order to have an independent inhibition, we substitute the optimal inhibition $WW^T$ with the correlation matrix of neural activity $\mathbb{E}[\mathbf{y}\mathbf{y}^T]$, yielding the substituted energy in equation 9. Taking the gradient of equation 9 with respect to the weights $W$ yields an update rule (Eqn. 10) which will drive the weights to grow without bound. This problem is shared by the original objective (Eqn. 4), which is also prone to weight explosion. We derive different rules for updating $W$ with learning rate $\eta$:

$$\text{(predictive)}\;\; W \leftarrow W + \eta(\mathbf{y} - W\mathbf{x})\mathbf{x}^\top, \qquad \text{(Hebbian)}\;\; W \leftarrow W + \eta(\mathbf{y}\mathbf{x}^\top - W). \tag{S2}$$

On whitened data $\mathbb{E}[W\mathbf{x}\mathbf{x}^\top] = W$, both rules have the same average update and hence the same fixed point (average update zero): $W^* = \mathbb{E}[\mathbf{y}\mathbf{x}^\top]$.

**Weight normalization.** Interestingly, we find in experiments that for more complex tasks (such as learning the non-Gaussian components on natural images), these update rules often choose to satisfy the weight penalty instead of the reconstruction objective, leading to weights being zero or very sparse. This is because the normalization penalty introduced in the substituted energies (Eqn. 11 and Eqn. 13) only prevents the weights from exploding, but it does not prevent them from shrinking towards zero. Therefore, we follow Olshausen and Field [1997] and constrain each neuron's feed-forward weight vector to be of unit norm. In the brain a candidate mechanism is synaptic scaling [Turrigiano et al., 1998], which has been proposed in models of feature learning before [Savin et al., 2010, Perrinet, 2010, Mayzel and Schneidman, 2024]. It is also worth noting that changing this norm will directly impact the consistency constraint, since the weights cannot change norms now, therefore we carefully choose the norm to be 0.1 which minimizes the consistency error for across rules.

**Input centering.** In neurons firing-rates are strictly non-negative, therefore, assuming most inputs come from other neurons, the input to the network will always have a strictly non-negative mean. This changes the construction of the consistency condition, which is based upon whitened centered data. With a positive mean, the condition that needs to be enforced is $\mathbf{y} \approx W(\mathbf{x} - \mu_{\mathbf{x}})$, which then allow us to write

$$WW^\top = W\mathbb{E}[(\mathbf{x} - \mu_{\mathbf{x}})(\mathbf{x} - \mu_{\mathbf{x}})^\top]W^\top = \mathbb{E}[W(\mathbf{x} - \mu_{\mathbf{x}})(\mathbf{x} - \mu_{\mathbf{x}})^\top W^\top] = \mathbb{E}[\mathbf{y}\mathbf{y}^\top]. \tag{S3}$$

Therefore, centering the input mostly tends to reduce the consistency error across all rules and datasets (Fig. S4A). Interestingly, reconstruction seems to be lower for non-centered networks that use the predictive or BCM rule (Fig. S4B). For the MNIST and images datasets, centering leads the same update rule to learn different receptive fields as we see in figures S5 and S6.

**Input normalization.** In order to keep the distribution of different datasets within the same range, we normalize the input by dividing it by its norm and multiplying by 3, such that $\mathbf{x}_i = \frac{\mathbf{x}_i}{3||\mathbf{x}_i||}$. This process is also biologically plausible, often termed divisive normalization [Heeger, 1992].

**Weight rectification.** The weight space can be constrained in multiple ways, but here we follow Dale's principle and hold all weights non-negative [Dale, 1935].

**Equivalence of objectives** In the following propositions, we show that the original predictive coding objective defined in equation 4 is equivalent to the predictive objective (Eqn. 13) up to atom length. Due to the scale degeneracy of the original objective (proposition 1.1), any implementation has to prevent the weights from exploding or shrinking towards zero and therefore, the magnitude of the weights is often kept constant throughout training. If the weights are kept within a consistency manifold, one can then learn the inhibition independently and minimize the simpler substituted energy (Eqn. 9), making the model more efficient and biologically plausible (proposition 1.2). Adding an input normalization term to the substituted energy will not affect the objective up to atom length (proposition 1.3). Instead, it will lead to to the consistency condition being optimally fulfilled (proposition 1.4), particularly when the predictive rule is applied (proposition 1.5).

**Consistency error** During the training of a network, we keep track of the distance between the weights $W$ and the consistency manifold, where $WW^\top = \mathbb{E}[\mathbf{y}\mathbf{y}^\top]$. This is termed the *consistency error* ($C$) and is defined as the normalized distance between $WW^\top$ and the network's estimate of the second moment $M$ (i.e. inhibition). On can write this as

$$C = \frac{||M - WW^T||_F}{||M||_F}. \tag{S4}$$

**Proposition 1.1** (The classical objective cannot fix atom norms)**.** *For spanning atoms, rescaling $W \to cW$, $\mathbf{y} \to \mathbf{y}/c$ leaves every reconstruction unchanged while sending the prior penalty to zero as $c \to \infty$. Every working algorithm therefore supplies a norm rule from outside the objective.*

*Proof.* We evaluate the objective $F$ with the rescaled dictionary $cW$. By definition,

$$F(cW) = \mathbb{E}\Big[\min_{\mathbf{y}}\big(\tfrac{1}{2}\|\mathbf{x} - (cW)^\top\mathbf{y}\|^2 + \phi(\mathbf{y})\big)\Big].$$

We perform a change of variables by defining $\mathbf{z} = c\mathbf{y}$. Since minimizing over $\mathbf{y}$ is equivalent to minimizing over $\mathbf{z}$, we can rewrite the objective as:

$$F(cW) = \mathbb{E}\Big[\min_{\mathbf{z}}\big(\tfrac{1}{2}\|\mathbf{x} - W^\top\mathbf{z}\|^2 + \phi(\mathbf{z}/c)\big)\Big].$$

The true minimum of this expression must be less than or equal to the value obtained by evaluating the inner function at any specific candidate $\mathbf{z}$. Because we assumed the $m$ rows of $W$ span $\mathbb{R}^n$ (where $m \geq n$), there exists at least one choice of $\mathbf{z}$ for every $\mathbf{x}$ that achieves perfect reconstruction, such that $W^\top\mathbf{z} = \mathbf{x}$.

By choosing this specific candidate $\mathbf{z}$ for each input, the reconstruction term $\frac{1}{2}\|\mathbf{x} - W^\top\mathbf{z}\|^2$ becomes exactly zero. Therefore, we establish an upper bound

$$F(cW) \leq \mathbb{E}[\phi(\mathbf{z}/c)].$$

Now, consider the limit as the scaling factor $c \to \infty$. For a fixed $\mathbf{z}$, $\mathbf{z}/c$ approaches the zero vector. Because the prior $\phi$ is continuous and normalized such that $\phi(\mathbf{0}) = 0$, we have $\phi(\mathbf{z}/c) \to 0$ as $c \to \infty$. Consequently, the expectation goes to zero, yielding $F(cW) \to 0$. In this limit, the codes $\mathbf{y} = \mathbf{z}/c$ also shrink to zero, therefore collapsing the model and yielding a trivial solution. □

**Proposition 1.2** (Energies are equivalent on the consistency manifold)**.** *For every $(\mathbf{y}, \mathbf{x})$ and every $W$,*

$$\tilde{E}(\mathbf{y}, \mathbf{x}) - E_P(\mathbf{y}, \mathbf{x}) = \tfrac{1}{2}\,\mathbf{y}^\top\big(WW^\top - \Sigma_y\big)\mathbf{y} \;+\; \tfrac{1}{2}\|\mathbf{x}\|^2. \tag{S5}$$

*The last term contains neither the code nor a code-dependent quantity, so it shifts every candidate code by the same amount and affects no minimizer. Hence the two energies induce identical codes exactly when $\Sigma_y = WW^\top$ (*consistency*); off the manifold, the codes differ in proportion to the gap $\Sigma_y - WW^\top$.*

*Proof.* We fully expand both energy functions and then take their difference.

First, consider the extended energy $\tilde{E} = E + \frac{1}{2}\|W\mathbf{x}\|^2$. Substituting the definition of $E$ from equation S1 and expanding the squared $L_2$ norm $\|\mathbf{x} - W^\top\mathbf{y}\|^2$:

$$\begin{aligned}
\tilde{E} &= \tfrac{1}{2}\|\mathbf{x} - W^\top\mathbf{y}\|^2 + \phi(\mathbf{y}) + \tfrac{1}{2}\|W\mathbf{x}\|^2 \\
&= \tfrac{1}{2}\big(\mathbf{x}^\top\mathbf{x} - 2\mathbf{y}^\top W\mathbf{x} + \mathbf{y}^\top WW^\top\mathbf{y}\big) + \phi(\mathbf{y}) + \tfrac{1}{2}\|W\mathbf{x}\|^2 \\
&= \tfrac{1}{2}\|\mathbf{x}\|^2 - \mathbf{y}^\top W\mathbf{x} + \tfrac{1}{2}\mathbf{y}^\top WW^\top\mathbf{y} + \tfrac{1}{2}\|W\mathbf{x}\|^2 + \phi(\mathbf{y}).
\end{aligned}$$

Next, consider the predictive energy $E_P$ from equation 13. Expanding the term $\frac{1}{2}\|\mathbf{y} - W\mathbf{x}\|^2$:

$$\begin{aligned} E_P &= \tfrac{1}{2}\|\mathbf{y} - W\mathbf{x}\|^2 + \tfrac{1}{2}\mathbf{y}^\top(\Sigma_y - I)\mathbf{y} + \phi(\mathbf{y}) \\ &= \tfrac{1}{2}\big(\mathbf{y}^\top\mathbf{y} - 2\mathbf{y}^\top W\mathbf{x} + (W\mathbf{x})^\top(W\mathbf{x})\big) + \tfrac{1}{2}\mathbf{y}^\top\Sigma_y\mathbf{y} - \tfrac{1}{2}\mathbf{y}^\top\mathbf{y} + \phi(\mathbf{y}) \\ &= \tfrac{1}{2}\|\mathbf{y}\|^2 - \mathbf{y}^\top W\mathbf{x} + \tfrac{1}{2}\|W\mathbf{x}\|^2 + \tfrac{1}{2}\mathbf{y}^\top\Sigma_y\mathbf{y} - \tfrac{1}{2}\|\mathbf{y}\|^2 + \phi(\mathbf{y}). \end{aligned}$$

Notice that the $+\frac{1}{2}\|\mathbf{y}\|^2$ and $-\frac{1}{2}\|\mathbf{y}\|^2$ terms cancel out, simplifying $E_P$ to:

$$E_P = -\mathbf{y}^\top W\mathbf{x} + \tfrac{1}{2}\|W\mathbf{x}\|^2 + \tfrac{1}{2}\mathbf{y}^\top\Sigma_y\mathbf{y} + \phi(\mathbf{y}).$$

Finally, we subtract $E_P$ from $\tilde{E}$. The shared terms $-\mathbf{y}^\top W\mathbf{x}$, $\frac{1}{2}\|W\mathbf{x}\|^2$, and $\phi(\mathbf{y})$ all cancel perfectly:

$$\begin{aligned} \tilde{E} - E_P &= \Big(\tfrac{1}{2}\|\mathbf{x}\|^2 + \tfrac{1}{2}\mathbf{y}^\top WW^\top\mathbf{y}\Big) - \Big(\tfrac{1}{2}\mathbf{y}^\top\Sigma_y\mathbf{y}\Big) \\ &= \tfrac{1}{2}\mathbf{y}^\top(WW^\top - \Sigma_y)\mathbf{y} + \tfrac{1}{2}\|\mathbf{x}\|^2. \end{aligned}$$

This completes the derivation of equation S5. □

**Proposition 1.3** (The added input normalization term does not change the solution)**.** *Let $\tilde{E} = E + \frac{1}{2}\|W\mathbf{x}\|^2$ and $\tilde{F}, F$ the corresponding objectives. Then: (a) latent code inference is unchanged, $\arg\min_{\mathbf{y}} \tilde{E} = \arg\min_{\mathbf{y}} E$, since the added term contains no $\mathbf{y}$; (b) on whitened data $\tilde{F}(W) = F(W) + \frac{1}{2}\|W\|_F^2$; and (c) every minimizer $W^\circ$ of $\tilde{F}$ minimizes the classical objective $F$ among all dictionaries of equal or smaller norm: $\|W'\|_F \leq \|W^\circ\|_F$ implies $F(W') \geq F(W^\circ)$.*

*Proof.* (a) The modified energy is $\tilde{E}(\mathbf{y}, \mathbf{x}) = E(\mathbf{y}, \mathbf{x}) + \frac{1}{2}\|W\mathbf{x}\|^2$. Since the added term $\frac{1}{2}\|W\mathbf{x}\|^2$ does not depend on the code $\mathbf{y}$, it acts as a constant during the minimization over $\mathbf{y}$. Adding a constant does not change the location of the minimum, hence $\arg\min_{\mathbf{y}} \tilde{E} = \arg\min_{\mathbf{y}} E$.

(b) We evaluate the expected value of the added term over the whitened data distribution. Using cyclic property of the trace operator and the fact that the data is whitened ($\mathbb{E}[\mathbf{x}\mathbf{x}^\top] = I$):

$$\begin{aligned} \mathbb{E}\big[\tfrac{1}{2}\|W\mathbf{x}\|^2\big] &= \tfrac{1}{2}\mathbb{E}\big[\mathbf{x}^\top W^\top W\mathbf{x}\big] \\ &= \tfrac{1}{2}\mathbb{E}\big[\mathrm{Tr}(\mathbf{x}^\top W^\top W\mathbf{x})\big] \\ &= \tfrac{1}{2}\mathrm{Tr}\big(W^\top W\mathbb{E}[\mathbf{x}\mathbf{x}^\top]\big) \\ &= \tfrac{1}{2}\mathrm{Tr}(W^\top W I) \\ &= \tfrac{1}{2}\|W\|_F^2. \end{aligned}$$

Therefore, taking the expectation of the minimum of $\tilde{E}$ yields the modified objective $\tilde{F}(W) = F(W) + \frac{1}{2}\|W\|_F^2$.

(c) We proof by contradiction. Let $W^\circ$ be a minimizer of the modified objective $\tilde{F}$. Suppose there exists another dictionary $W'$ that achieves a strictly better classical objective $F(W') < F(W^\circ)$, while having an equal or smaller norm, meaning $\|W'\|_F \leq \|W^\circ\|_F$.

Using the identity from part (b), we evaluate $\tilde{F}$ for $W'$:

$$\tilde{F}(W') = F(W') + \tfrac{1}{2}\|W'\|_F^2.$$

Applying our assumptions $F(W') < F(W^\circ)$ and $\frac{1}{2}\|W'\|_F^2 \leq \frac{1}{2}\|W^\circ\|_F^2$, we obtain:

$$\tilde{F}(W') < F(W^\circ) + \tfrac{1}{2}\|W^\circ\|_F^2 = \tilde{F}(W^\circ).$$

This implies $\tilde{F}(W') < \tilde{F}(W^\circ)$, which directly contradicts the premise that $W^\circ$ is a minimizer of $\tilde{F}$. Thus, no such $W'$ can exist. □

**Proposition 1.4** (Residual)**.** *Define $\mathbf{e} = \mathbf{y} - W^*\mathbf{x}$, the part of the code not predicted by $W^*$, with covariance $\Sigma_e = \mathbb{E}[\mathbf{e}\mathbf{e}^\top]$. Then $\mathbb{E}[\mathbf{e}\mathbf{x}^\top] = 0$ and*

$$\Sigma_y = W^*W^{*\top} + \Sigma_e : \tag{S6}$$

*the fixed point sits on the consistency manifold up to exactly $\Sigma_e$. For the Gaussian prior, $\mathbf{e} = 0$ on every sample.*

*Proof.* First, we show that the residual $\mathbf{e}$ is uncorrelated with the input $\mathbf{x}$. By definition, $\mathbf{e} = \mathbf{y} - W^*\mathbf{x}$. Taking the expectation of the outer product with $\mathbf{x}$:

$$\begin{aligned}\mathbb{E}[\mathbf{e}\mathbf{x}^\top] &= \mathbb{E}[(\mathbf{y} - W^*\mathbf{x})\mathbf{x}^\top] \\ &= \mathbb{E}[\mathbf{y}\mathbf{x}^\top] - W^*\mathbb{E}[\mathbf{x}\mathbf{x}^\top].\end{aligned}$$

Recall the fixed point of the learning rules is $W^* = \mathbb{E}[\mathbf{y}\mathbf{x}^\top]$, and the data is whitened such that $\mathbb{E}[\mathbf{x}\mathbf{x}^\top] = I$. Substituting these yields:

$$\mathbb{E}[\mathbf{e}\mathbf{x}^\top] = W^* - W^*I = \mathbf{0}.$$

Next, we derive the covariance of the code $\mathbf{y}$, denoted as $\Sigma_y$. We substitute $\mathbf{y} = W^*\mathbf{x} + \mathbf{e}$ into the covariance definition:

$$\begin{aligned}\Sigma_y &= \mathbb{E}[\mathbf{y}\mathbf{y}^\top] \\ &= \mathbb{E}[(W^*\mathbf{x} + \mathbf{e})(W^*\mathbf{x} + \mathbf{e})^\top] \\ &= \mathbb{E}[W^*\mathbf{x}\mathbf{x}^\top W^{*\top} + W^*\mathbf{x}\mathbf{e}^\top + \mathbf{e}\mathbf{x}^\top W^{*\top} + \mathbf{e}\mathbf{e}^\top].\end{aligned}$$

Using linearity of expectation, we distribute the expected value. Since $\mathbb{E}[\mathbf{e}\mathbf{x}^\top] = 0$, its transpose $\mathbb{E}[\mathbf{x}\mathbf{e}^\top]$ is also 0. Thus, the cross-terms vanish:

$$\begin{aligned}\Sigma_y &= W^*\mathbb{E}[\mathbf{x}\mathbf{x}^\top]W^{*\top} + W^*\mathbb{E}[\mathbf{x}\mathbf{e}^\top] + \mathbb{E}[\mathbf{e}\mathbf{x}^\top]W^{*\top} + \mathbb{E}[\mathbf{e}\mathbf{e}^\top] \\ &= W^*IW^{*\top} + 0 + 0 + \Sigma_e \\ &= W^*W^{*\top} + \Sigma_e.\end{aligned}$$

This establishes equation S6.

Finally, in the case of a Gaussian prior, the prior penalty $\phi(\mathbf{y})$ is a quadratic function of $\mathbf{y}$. This makes the entire substituted energy $E_P(\mathbf{y}, \mathbf{x})$ a purely quadratic form. The minimizer of a strictly quadratic objective with respect to $\mathbf{x}$ is a linear projection. Therefore, the optimal code takes the exact linear form $\mathbf{y} = B\mathbf{x}$ for some matrix $B$.

Computing the fixed point under this strictly linear relationship gives

$$W^* = \mathbb{E}[\mathbf{y}\mathbf{x}^\top] = \mathbb{E}[B\mathbf{x}\mathbf{x}^\top] = B\mathbb{E}[\mathbf{x}\mathbf{x}^\top] = B.$$

Since $W^* = B$, the code is exactly $\mathbf{y} = W^*\mathbf{x}$. The residual is therefore $\mathbf{e} = \mathbf{y} - W^*\mathbf{x} = \mathbf{0}$ for every sample, which means $\Sigma_e = \mathbf{0}$. ☐

**Proposition 1.5** (Error dynamics and the leftover kick)**.** *Let $\boldsymbol{\delta}_t = W_t - W^*$ denote the parameter error. For both learning rules, the exact update can be decomposed into a shared contraction, a baseline "kick" $\boldsymbol{\xi}_t$, and a higher-order interaction term. The kick $\boldsymbol{\xi}_t$ is defined as the exact update that remains even if the weights are already correct ($\boldsymbol{\delta}_t = 0$):*

$$\boldsymbol{\delta}_{t+1} \approx \underbrace{\boldsymbol{\delta}_t\big(I - \eta\,\mathbf{x}_t\mathbf{x}_t^\top\big)}_{\text{shared contraction}} + \eta \underbrace{\boldsymbol{\xi}_t}_{\text{leftover kick}}, \qquad \boldsymbol{\xi}_t = \begin{cases}\mathbf{e}_t\mathbf{x}_t^\top & \textit{(predictive)},\\ \mathbf{e}_t\mathbf{x}_t^\top + W^*(\mathbf{x}_t\mathbf{x}_t^\top - I) & \textit{(Hebbian)}.\end{cases} \tag{S7}$$

*Because the Hebbian kick contains the persistent noise term $W^*(\mathbf{x}_t\mathbf{x}_t^\top - I)$, it pushes the weights off the consistency manifold on every sample. By Proposition 1.2, moving off this manifold causes the Hebbian network to infer incorrect codes, creating a feedback loop that permanently separates its trajectory from optimal learning.*

*Proof.* We find the exact error dynamics by substituting $\mathbf{y}_t = W^*\mathbf{x}_t + \mathbf{e}_t$ and $W_t = W^* + \boldsymbol{\delta}_t$ into the parameter updates. For the predictive rule, the substitution is

$$\begin{aligned}W_{t+1} &= W_t + \eta(\mathbf{y}_t - W_t\mathbf{x}_t)\mathbf{x}_t^\top \\ W^* + \boldsymbol{\delta}_{t+1} &= W^* + \boldsymbol{\delta}_t + \eta\big(W^*\mathbf{x}_t + \mathbf{e}_t - (W^* + \boldsymbol{\delta}_t)\mathbf{x}_t\big)\mathbf{x}_t^\top \\ \boldsymbol{\delta}_{t+1} &= \boldsymbol{\delta}_t - \eta\boldsymbol{\delta}_t\mathbf{x}_t\mathbf{x}_t^\top + \eta\mathbf{e}_t\mathbf{x}_t^\top \\ \boldsymbol{\delta}_{t+1} &= \underbrace{\boldsymbol{\delta}_t\big(I - \eta\mathbf{x}_t\mathbf{x}_t^\top\big)}_{\text{shared contraction}} + \eta \underbrace{\big(\mathbf{e}_t\mathbf{x}_t^\top\big)}_{\text{predictive kick}}.\end{aligned}$$

For the Hebbian rule, substituting the same terms yields

$$\begin{aligned}W_{t+1} &= W_t + \eta(\mathbf{y}_t\mathbf{x}_t^\top - W_t) \\ W^* + \boldsymbol{\delta}_{t+1} &= W^* + \boldsymbol{\delta}_t + \eta\big((W^*\mathbf{x}_t + \mathbf{e}_t)\mathbf{x}_t^\top - (W^* + \boldsymbol{\delta}_t)\big) \\ \boldsymbol{\delta}_{t+1} &= \boldsymbol{\delta}_t + \eta W^*\mathbf{x}_t\mathbf{x}_t^\top + \eta\mathbf{e}_t\mathbf{x}_t^\top - \eta W^* - \eta\boldsymbol{\delta}_t.\end{aligned}$$

To isolate the shared contraction term $\boldsymbol{\delta}_t(I - \eta \mathbf{x}_t\mathbf{x}_t^\top)$, we rewrite $-\eta\boldsymbol{\delta}_t$ by adding and subtracting $\eta\boldsymbol{\delta}_t\mathbf{x}_t\mathbf{x}_t^\top$. This allows us to regroup the exact equation into three distinct parts as

$$\boldsymbol{\delta}_{t+1} = \underbrace{\boldsymbol{\delta}_t\big(I - \eta\mathbf{x}_t\mathbf{x}_t^\top\big)}_{\text{shared contraction}} + \eta\underbrace{\big(\mathbf{e}_t\mathbf{x}_t^\top + W^*(\mathbf{x}_t\mathbf{x}_t^\top - I)\big)}_{\text{baseline kick evaluated at } \boldsymbol{\delta}_t=0} + \underbrace{\eta\boldsymbol{\delta}_t(\mathbf{x}_t\mathbf{x}_t^\top - I)}_{\text{noise interacting with error}}.$$

The first term is the *shared contraction*. Because the input data is whitened ($\mathbb{E}[\mathbf{x}\mathbf{x}^\top] = I$), the expectation of this term is $(1-\eta)\boldsymbol{\delta}_t$. This provides a deterministic pull that reduces the error toward zero on average.

The second term, the *leftover kick* $\boldsymbol{\xi}_t$, is the baseline noise injected into the system even if the weights are perfectly correct ($\boldsymbol{\delta}_t = 0$). While this kick averages to zero in expectation for both rules (ensuring they share the same fixed point $W^*$), on every individual sample, the Hebbian extra term $W^*(\mathbf{x}_t\mathbf{x}_t^\top - I)\big)$ pushes the weights off the consistency manifold. Because the computation of the correct code requires on-manifold energy, the Hebbian rule leads to incorrect inference and non-optimal updates.

The third term, $\eta\boldsymbol{\delta}_t(\mathbf{x}_t\mathbf{x}_t^\top - I)$, is the interaction between the sampling noise and the current parameter error. As the network learns and $\boldsymbol{\delta}_t$ approaches zero, this term vanishes entirely. □

#### S2 Inference dynamics for different priors

Here we describe the activity (i.e. inference) dynamics that emerge for different choices of priors $\phi$. The weight gradients do not depend on the prior and therefore remain the same for all priors.

##### Gaussian prior

We re-write equation 1 with an explicit noise variance $\sigma_\varepsilon^2$ as

$$\mathbf{x} = D\mathbf{y} + \varepsilon, \qquad \varepsilon \sim \mathcal{N}(0, \sigma_\varepsilon^2 I). \tag{S8}$$

Setting the prior to be Gaussian with mean 0 and covariance $\sigma_y^2 I$ yields $\phi(\mathbf{y}) = \frac{c}{2}||\mathbf{y}||_2^2$, where $c = \sigma_\epsilon^2/\sigma_y^2$ is the ratio of the observation noise variance to the prior variance (energies are in units of $\sigma_\epsilon^2$). This yields the energy function with input normalization (cf. Eqn. 13):

$$E_G(\mathbf{y}, \mathbf{x}; W) = \frac{1}{2}||\mathbf{x}||_2^2 - \mathbf{x}^T W^T \mathbf{y} + \frac{1}{2}\mathbf{y}^T M \mathbf{y} + \frac{1}{2}||W\mathbf{x}||_2^2 + \frac{c}{2}||\mathbf{y}||_2^2 \tag{S9}$$

For the inference step, the gradient with respect to $\mathbf{y}$ is

$$\nabla_{\mathbf{y}} E_G = -W\mathbf{x} + M\mathbf{y} + c\,\mathbf{y}. \tag{S10}$$

Finding the minimum of this energy function yields

$$\mathbf{y}^* = (M + cI)^{-1} W\mathbf{x}. \tag{S11}$$

Inference can be performed by a linear network (with step size $\tau$), which runs until a stable state is found:

$$\mathbf{y}_{t+1} = \mathbf{y}_t + \tau(W\mathbf{x} - M\mathbf{y}_t - c\,\mathbf{y}_t) \tag{S12}$$

The term $c\,\mathbf{y}_t$ is a leak with size determined by the prior. A broad prior relative to the observation noise gives a weak leak, and equal variances give a leak of unity. The consistency conditions require $\mathbf{y} = W\mathbf{x}$, so it requires $M + cI = I$. Therefore, this requires $c < 1$, or $\sigma_y^2 > \sigma_\epsilon^2$, to hold, i.e. the latent variance must exceed the observation noise.

##### Exponential prior

Setting the prior to be exponential $\phi(\mathbf{y}) = \lambda\mathbf{1}^T\mathbf{y}$ with $\mathbf{y} \geq 0$, with rate $\lambda$ restricts the latent variables to be non-negative. The corresponding energy function is

$$E_E(\mathbf{y}, \mathbf{x}; W) = \frac{1}{2}||\mathbf{x}||_2^2 - \mathbf{x}^T W^T \mathbf{y} + \frac{1}{2}\mathbf{y}^T M \mathbf{y} + \frac{1}{2}||W\mathbf{x}||_2^2 + \lambda\mathbf{1}^T\mathbf{y}, \quad \mathbf{y} \geq 0. \tag{S13}$$

On the non-negative part of the function, the gradient with respect to $\mathbf{y}$ is

$$\nabla_{\mathbf{y}} E_E = -W\mathbf{x} + M\mathbf{y} + \lambda \mathbf{1}. \tag{S14}$$

Unlike the Gaussian case (Eqn. S10), the exponential prior contributes a constant term $\lambda$ instead of a leak. In this case, stability is obtained through the positive diagonal of $M$, and not by by the prior. This can be seen when considering the requirement $\mathbf{y} = W\mathbf{x}$, where consistency is satisfied at $M = WW^\top = I$. Therefore, for exponential priors both consistency conditions can be met, in principle, exactly. However, rectification limits the range of $\mathbf{y}$, so may sparsify the encoding, and for an overcomplete representation $WW^\top$ has rank at most $n < m$, so does not fully determine $M$.

Setting the gradient to 0 does not yield a closed-form solution due to the non-negativity constraint must be enforced. Instead, following Rozell et al. [2008] inference can instead be performed via projected (proximal) gradient descent, where each gradient step is followed by a projection onto the non-negative space, by

$$\mathbf{y}_{t+1} = \left[\mathbf{y}_t + \tau(W\mathbf{x} - M\mathbf{y}_t - \lambda\mathbf{1})\right]_+ . \tag{S15}$$

Here, $[\cdot]_+ = \max(0, \cdot)$ is element-wise rectification to ensure non-negative responses.

##### Laplace prior

Setting the prior a Laplace distribution models the latent variables as sparse, regardless of their sign, so it does not meet the biological constraint of non-negativity. We include it as it was interdicted as a sparseness constraint in sparse coding models [Olshausen and Field, 1997]. This corresponds to setting $\phi(\mathbf{y}) = \lambda||\mathbf{y}||_1$, with an energy function

$$E_L(\mathbf{y}, \mathbf{x}; W) = \frac{1}{2}||\mathbf{x}||_2^2 - \mathbf{x}^T W^T \mathbf{y} + \frac{1}{2}\mathbf{y}^T M \mathbf{y} + \frac{1}{2}||W\mathbf{x}||_2^2 + \lambda||\mathbf{y}||_1. \tag{S16}$$

Because the $L_1$ norm is not differentiable at zero, we consider the sub-gradient of the energy

$$\partial_{\mathbf{y}} E_L = -W\mathbf{x} + M\mathbf{y} + \lambda \text{sign}(\mathbf{y}). \tag{S17}$$

This shows that the Laplace prior contributes a piecewise constant term $\lambda\,\text{sign}(\mathbf{y})$ to the gradient. This means that, as for the exponential case, the prior does not contribute a leak term. Additionally, the Laplace prior does not restrict the sign of the latent variables, so the network can be consistent with $M = WW^\top$ and $\mathbf{y} = W\mathbf{x}$. On the other hand, the rank constraint for an over-complete representation still applies.

Similar to the exponential prior, setting the sub-gradient to 0 does not yield a simple closed-form linear solution. Instead, inference is commonly performed using the Iterative Shrinkage-Thresholding Algorithm (ISTA) [Daubechies et al., 2004], which applies a gradient step on the smooth part of the energy function followed by a proximal operator for the $L_1$ penalty

$$\mathbf{y}_{t+1} = \mathcal{S}_{\lambda\tau}\left(\mathbf{y}_t + \tau(W\mathbf{x} - M\mathbf{y}_t)\right). \tag{S18}$$

Here, $\mathcal{S}_\alpha(v) = \text{sign}(v)\max(0, |v| - \alpha)$ is the element-wise soft-thresholding operator. This function shrinks small activities to exactly zero, inducing a sparse representation without restricting the network to strictly non-negative values.

#### S3 Plasticity rules

In this work, we compare a variety of local and biologically plausible plasticity rules.

##### Hebbian rule

A biologically plausible rule can be obtained by computing the gradient of the substituted energy with weight normalization (equation 11), obtaining

$$\nabla_{\mathbf{y}} E_H = -\mathbf{y}\mathbf{x}^\top + W. \tag{S19}$$

The update is then computed by performing gradient descent on this gradient, obtaining the rule specified in equation S19. In order to obtain the fixed point, we can set $\mathbb{E}[\Delta W] = 0$ and obtain:

$$
\begin{aligned}
0 &= \mathbb{E}[-\mathbf{y}\mathbf{x}^\top + W] \\
&= -\mathbb{E}[\mathbf{y}\mathbf{x}^\top] + W
\end{aligned}
\tag{S20}
$$

Therefore, the feed-forward weight matrix at the fixed point is

$$
W^* = \mathbb{E}[\mathbf{y}\mathbf{x}^\top]. \tag{S21}
$$

**Oja rule**

The Hebbian rule requires an explicit weight decay term to prevent unbounded growth. Oja's rule [Oja, 1982] achieves normalization implicitly by coupling the decay to the postsynaptic activity of each neuron individually. In its original form, each output unit $i$ with response $y_i = \mathbf{w}_i^T \mathbf{x}$ updates its weight vector as $\Delta \mathbf{w}_i = \eta\, y_i(\mathbf{x} - y_i \mathbf{w}_i)$. Stacking these rows gives the full-matrix update

$$
W \leftarrow W + \eta \left( \mathbf{y}\mathbf{x}^T - \mathrm{diag}(\mathbf{y} \odot \mathbf{y})\, W \right). \tag{S22}
$$

Here, $\odot$ is the element-wise product and $\mathrm{diag}(\mathbf{y} \odot \mathbf{y})$ is the diagonal matrix of squared responses $y_i^2$. Since the gradient we are optimizing here is $\Delta W = \mathbf{y}\mathbf{x}^T - \mathrm{diag}(\mathbf{y} \odot \mathbf{y})\, W$, we can set $\mathbb{E}[\Delta W] = 0$ and obtain, for each row $i$

$$
\begin{aligned}
0 &= \mathbb{E}[y_i \mathbf{x}^T - y_i^2 \mathbf{w}_i^T] \\
&= \mathbb{E}[y_i \mathbf{x}^T] - \mathbb{E}[y_i^2]\, \mathbf{w}_i^T.
\end{aligned}
$$

Therefore, each row of the feed-forward weight matrix at the fixed point is

$$
\mathbf{w}_i = \frac{\mathbb{E}[y_i \mathbf{x}]}{\mathbb{E}[y_i^2]}, \qquad \text{equivalently} \qquad W^* = \mathrm{diag}(\mathbb{E}[\mathbf{y} \odot \mathbf{y}])^{-1} \mathbb{E}[\mathbf{y}\mathbf{x}^T]. \tag{S23}
$$

**Földiák rule**

Another plausible feed-forward rule proposed in the literature is the Földiák rule [Földiak, 1990], in which the decay term is gated by the postsynaptic activity and drives each weight toward the input rather than toward zero. Per synapse, the rule is defined as $\Delta w_{ij} = \eta\, y_i(x_j - w_{ij})$, which in matrix form is

$$
W \leftarrow W + \eta\, \mathrm{diag}(\mathbf{y}) \left( \mathbf{1}\mathbf{x}^T - W \right), \tag{S24}
$$

where $\mathbf{1}\mathbf{x}^T$ is the matrix whose every row equals $\mathbf{x}^T$. Since the gradient we are optimizing is $\Delta W = \mathrm{diag}(\mathbf{y})(\mathbf{1}\mathbf{x}^T - W)$, we can set $\mathbb{E}[\Delta W] = 0$ and obtain, for each entry $(i, j)$,

$$
\begin{aligned}
0 &= \mathbb{E}[y_i(x_j - w_{ij})] \\
&= \mathbb{E}[y_i x_j] - \mathbb{E}[y_i]\, w_{ij}.
\end{aligned}
$$

Therefore, the feed-forward weight matrix at the fixed point is

$$
W^* = \mathrm{diag}(\mathbb{E}[\mathbf{y}])^{-1} \mathbb{E}[\mathbf{y}\mathbf{x}^T]. \tag{S25}
$$

**BCM-like rule**

The Bienenstock–Cooper–Munro (BCM) rule [Bienenstock et al., 1982] modulates plasticity through a sliding threshold $\boldsymbol{\theta}$ that sets the sign of the weight change. We use the simplified update

$$
W_t = W_{t-1} + \eta \left( g(\mathbf{y}, \boldsymbol{\theta}) \mathbf{x}^T - W_{t-1} \right), \tag{S26}
$$

and define

$$g(\mathbf{y}, \boldsymbol{\theta}) = \mathbf{y} - \boldsymbol{\theta}, \qquad \boldsymbol{\theta} = \mathbb{E}[\mathbf{y}]. \tag{S27}$$

Here the threshold is the mean activity rather than the second moment $\mathbb{E}[y_i^2]$ of the standard rule. Since the gradient we are optimizing is $\Delta W = (\mathbf{y} - \boldsymbol{\theta})\mathbf{x}^T - W$, we can set $\mathbb{E}[\Delta W] = 0$ and obtain

$$\begin{aligned} 0 &= \mathbb{E}[(\mathbf{y} - \boldsymbol{\theta})\mathbf{x}^T - W] \\ &= \mathbb{E}[\mathbf{y}\mathbf{x}^T] - \boldsymbol{\theta}\,\mathbb{E}[\mathbf{x}]^T - W. \end{aligned}$$

Substituting the threshold with mean activity, we obtain the feed-forward weight matrix at the fixed point

$$W^* = \mathbb{E}[\mathbf{y}\mathbf{x}^T] - \mathbb{E}[\mathbf{y}]\,\mathbb{E}[\mathbf{x}]^T. \tag{S28}$$

**Predictive rule**

The rule we introduce in this work can be directly obtained by computing the gradient of the substituted energy with input normalization (equation 13), obtaining

$$\nabla_{\mathbf{y}} E_P = -(\mathbf{y}\mathbf{x}^\top + W\mathbf{x}\mathbf{x}^\top). \tag{S29}$$

The update is then computed by performing gradient descent on this gradient, obtaining the rule specified in equation 14. In order to obtain the fixed point, we can set $\mathbb{E}[\Delta W] = 0$ and obtain

$$\begin{aligned} 0 &= \mathbb{E}[-\mathbf{y}\mathbf{x}^\top + W\mathbf{x}\mathbf{x}^\top] \\ &= -\mathbb{E}[\mathbf{y}\mathbf{x}^\top] + W\mathbb{E}[\mathbf{x}\mathbf{x}^\top]. \end{aligned} \tag{S30}$$

Therefore, the feed-forward weight matrix at the fixed point is

$$W^* = \mathbb{E}[\mathbf{y}\mathbf{x}^\top](\mathbb{E}[\mathbf{x}\mathbf{x}^\top])^{-1}. \tag{S31}$$

For whitened data, this is the same fixed point as the Hebbian rule.

#### S4 Consistency error for predictive rule and Gaussian prior

As the predictive rule follows the consistency manifold, and the model with Gaussian prior has a closed-form solution, we can compute the consistency error between the feed-forward weights and the recurrent weights at the fixed point. This requires the two fixed points of the network, both for inference and for learning.

Substituting the inference fixed point (Eqn. S11) into the inhibitory fixed point $M = \mathbb{E}[\mathbf{y}\mathbf{y}^\top]$ of Eqn. 8 gives $M = (M + cI)^{-1} W \Sigma_x W^\top (M + cI)^{-1}$, and multiplying by $M + cI$ on both sides yields

$$(M + cI)\, M\, (M + cI) = W \Sigma_x W^\top, \tag{S32}$$

where $\Sigma_x = \mathbb{E}[\mathbf{x}\mathbf{x}^\top]$ is the input second moment, or the covariance when the input is centered. As the left-hand side is a polynomial in $M$, the two sides can be compared mode by mode below.

Writing the feed-forward weights as $W = VS\tilde{U}^\top$ yields the output directions $V$, singular values $S = \mathrm{diag}(s_1, \ldots, s_r)$ (with rank $r = \mathrm{rank}\, W$), and encoded input directions $\tilde{U}$. We write $a_i := s_i^2$ for the eigenvalues of $WW^\top$. Then the encoded input becomes

$$W \Sigma_x W^\top = V\, SGS\, V^\top, \qquad G := \tilde{U}^\top \Sigma_x \tilde{U}, \tag{S33}$$

where $G$ is the input second moment restricted to the encoded subspace.

As the inhibition $M$ only acts on the feed-forward encoded subspace (cf. Eqn. S32), we can write $M = VNV^\top$ with $N \in \mathbb{R}^{r \times r}$ symmetric positive definite. Projecting Eqn. S32 with $V^\top(\cdot)V$ then gives

$$(N + cI)N(N + cI) = SGS. \tag{S34}$$

**On the consistency manifold.** The condition $M = WW^\top$ yields $VNV^\top = VS^2V^\top$, i.e. $N = S^2$. The left side of Eqn. S34 is then diagonal with entries $a_i(a_i + c)^2$, so

$$G = S^{-1}\operatorname{diag}\bigl(a_i(a_i + c)^2\bigr)S^{-1} = \operatorname{diag}\bigl((a_i + c)^2\bigr). \tag{S35}$$

Consistency therefore requires the encoded directions to be both orthonormal and mutually $\Sigma_x$-orthogonal, $\tilde{u}_i^\top \Sigma_x \tilde{u}_j = 0$ for $i \neq j$, and they are forced to be principal axes of $\Sigma_x$ when $r$ equals the input dimension. With

$$\nu_i := \tilde{u}_i^\top \Sigma_x \tilde{u}_i = G_{ii} = \mathbb{E}\bigl[(\tilde{u}_i^\top \mathbf{x})^2\bigr] \tag{S36}$$

for the input variance along encoded direction $i$ (an eigenvalue of $\Sigma_x$ in the aligned case, or in general a convex combination of eigenvalues), the gain on the manifold is

$$a_i = \sqrt{\nu_i} - c, \tag{S37}$$

which is positive for $\nu_i > c^2$ as directions carrying less variance than the squared leak cannot be encoded.

**At the fixed point of the predictive rule.** The second condition is given by the predictive learning rule. With $K = (M + cI)^{-1}$, the expected update of Eqn. 14 is $\mathbb{E}[\Delta W]/\eta = (K - I)W\Sigma_x$, so $\mathbb{E}[\Delta W] = 0$ gives $KW = W$, that is $MW = (1 - c)W$ and hence $N = (1 - c)I_r$. This requires $c < 1$, and it recovers condition (2) of Eqn. 7 as a property of the rule, since $M + cI = I$ on the encoded subspace and therefore $\mathbf{y} = W\mathbf{x}$ exactly. Substituting $N = (1 - c)I$ into Eqn. S34 gives $SGS = (1 - c)I$, so $\nu_i = (1 - c)/a_i$ and

$$a_i = \frac{1 - c}{\nu_i}. \tag{S38}$$

The code variance is then $\operatorname{Var}(y_i) = a_i\nu_i = 1 - c$, which is identical in every mode. Therefore, the learning rule whitens and assigns small gains to high-variance directions and large gains to low-variance ones, an opposite ordering to the principal subspace projection.

Eqns. S37 and S38 are decreasing and increasing function of $\nu_i$, respectively. Equating them gives $\nu_i^{3/2} - c\,\nu_i - (1-c) = 0$, with only positive root is $\nu_i = 1$, where both give $a_i = 1 - c$.

**The consistency error.** At the fixed point of the rule, $M = (1 - c)VV^\top$ and $WW^\top = V\operatorname{diag}\bigl((1 - c)/\nu_i\bigr)V^\top$, so

$$M - WW^\top = (1 - c)\,V\operatorname{diag}\bigl(1 - \nu_i^{-1}\bigr)V^\top. \tag{S39}$$

Since $V$ has orthonormal columns, $\|VDV^\top\|_F = \|D\|_F$ for any symmetric $D$, giving $\|M\|_F = (1 - c)\sqrt{r}$ and

$$\|M - WW^\top\|_F = (1 - c)\sqrt{\sum_{i=1}^{r}\bigl(1 - \nu_i^{-1}\bigr)^2}, \tag{S40}$$

with the sum running over encoded directions (we assume full rank of $W$ in our analysis). Dividing by $\|M\|_F$ cancels the factor $1 - c$, so one can write the consistency error $C$ (Eqn.S4) as a function of the input spectrum alone, independent of the prior variance and of the overall activity scale:

$$C = \sqrt{\frac{1}{r}\sum_{i=1}^{r}\bigl(1 - \nu_i^{-1}\bigr)^2} = \operatorname{RMS}_i\bigl|1 - \nu_i^{-1}\bigr|. \tag{S41}$$

This normalized consistency error $C$ is plotted in Fig. 2E. In the main text we use the eigenvalues of the input second moment $\Sigma_x$ to compute the consistency error from the data directly. In a misaligned network this is a conservative estimate of the error, as then the encoded directions are not aligned with the principal axes of $\Sigma_x$ and therefore $\nu_i$ is a convex combination of eigenvalues.

#### S5 Continuous excitatory-inhibitory model

##### Dynamics

Similar to the discrete/sequential model, the continuous excitatory-inhibitory model has a feed-forward matrix $\mathbf{W} \in \mathbb{R}^{n\times m}$ and a slow recurrent excitatory matrix $\mathbf{P} \in \mathbb{R}^{n\times n}$. In order to have two populations of neurons, we introduce a new population of inhibitory interneurons $\mathbf{z} \in \mathbb{R}^q$, excitatory to inhibitory weights $V \in \mathbb{R}^{q\times n}$, inhibitory to inhibitory weights $M^{q\times n}$ and inhibitory to excitatory weights $U \in \mathbb{R}^{n\times q}$ Instead of retrieving a static feedback signal, recurrent excitation now integrates past activity with a kernel $K$ which we describe below. Similar to the models analysed in the last chapter, we define the continuous model in terms of a discretized Euler simulation with step size $\eta_E$. Given a stream of data $\{\mathbf{x}_i\}_{i=1}^N$, the neural dynamics are updated according to the following equation:

$$\mathbf{y}_{t+1} = \mathbf{y}_t + \eta_E(\mathbf{W}\mathbf{x}_{t+1} + \mathbf{P}K(\mathbf{y}_t, ..., \mathbf{y}_{t-\beta}) - \mathbf{U}[\mathbf{z}_t]_+) \tag{S42}$$

$$\mathbf{z}_{t+1} = \mathbf{z}_t + \eta_E(\mathbf{V}\mathbf{y}_{t+1} - \mathbf{M}[\mathbf{z}_t]_+) \tag{S43}$$

##### Temporal kernel

The kernel function $K : \mathbb{R}^{n_y\beta} \to \mathbb{R}^{n_y}$ summarizes all the information from the past $\beta$ time steps. Let $x \in [0, 1]$, we use the alpha kernel due to its biological plausibility, denoted as $A(x; \mu, \sigma)$, is defined as:

$$A(x; \mu, \sigma) = \begin{cases} \frac{(x-\mu)}{\sigma^2} \exp\left(-\frac{(x-\mu)}{\sigma}\right), & \text{if } x \geq \mu, \\ 0, & \text{if } x < \mu, \end{cases} \tag{S44}$$

Here, $\mu$ is the location parameter and $\sigma > 0$ controls the spread and skewness of the function. Given the kernel function, we compute the recurrent excitatory input by integrating the past $\beta$ time steps with the kernel, such that:

$$K(\mathbf{y}_t, ..., \mathbf{y}_{t-\beta}) = \sum_{i=0}^{\beta} k(i/\beta)\mathbf{y}_{t-i} \tag{S45}$$

##### Asynchronous updates

We follow a similar strategy to [Aguiar and Hennig, 2024] to enable the models learn from a continuous stream of data. An important difference is that our plasticity threshold is now stochastic. Each time step a postsynaptic neuron updates its incoming synapses if its firing rate $v_i(t)$ and the time elapsed since its last update $\tau_i(t)$ simultaneously exceed thresholds $\theta_i^v(t)$ and $\theta_i^\tau(t)$ respectively. These thresholds are random variables drawn independently at every step, such that

$$\theta_i^v(t) \sim \mathcal{N}\big(\mu_v, \sigma_v^2\big), \qquad \theta_i^\tau(t) \sim \mathcal{N}\big(\mu_\tau, \sigma_\tau^2\big). \tag{S46}$$

Here, $\mu_v, \sigma_v$ and $\mu_\tau, \sigma_\tau$ are fixed parameters shared amongst all neurons. For the results in the main text, we set $\mu_v = 1.4, \sigma_v = 0.4, \mu_\tau = 10, \sigma_\tau = 1$ for a time constant of $\tau = 1$. The update for the post-synpatic neuron is gated according to

$$s_i(t) \sim \text{Bernoulli}(p_i(t)), \qquad p_i(t) = \mathbb{P}[\theta_i^v \leq v_i(t),\ \theta_i^\tau \leq \tau_i(t)] = \Phi(\tilde{v}_i(t))\ \Phi(\tilde{\tau}_i(t)), \tag{S47}$$

with $\tilde{v}_i(t) = (v_i(t) - \mu_v)/\sigma_v$, $\tilde{\tau}_i(t) = (\tau_i(t) - \mu_\tau)/\sigma_\tau$ and $\Phi$ the standard normal CDF. The factorisation follows from the independence of the two thresholds, so $p_i(t)$ is simply their joint CDF evaluated at the current state of the neuron; sampling a single Bernoulli variable is therefore equivalent to, but cheaper than, drawing both thresholds explicitly. Whenever $s_i(t) = 1$ the incoming weights of neuron $i$ are updated and its counter is reset, $\tau_i(t) \leftarrow 0$; otherwise the counter is incremented by the time constant $\tau_i(t) \leftarrow \tau_i(t-1) + \tau$.

Two properties of this rule are worth emphasising. First, a neuron that has updated recently has $\Phi(\tilde{\tau}_i) \approx 0$ and is no longer performing updates regardless of how strongly it is driven, which reproduces the effect of a refractory period without imposing a hard time window. Second, the deterministic rule of Aguiar and Hennig [2024] is recovered in the limit $\sigma_v, \sigma_\tau \to 0$, where the neuron updates if $v_i > \mu_v$ and $\tau_i > \mu_\tau$. This stochastic rule is important since it allows neurons to perform LTD updates when the activity low. If a deterministic threshold is set, then the mean activity of the neuron will revolve around that threshold, leading rules like the BCM and predictive to only apply LTP updates.

#### S6 Stimuli

We evaluate the model on stimuli ordered from fully controlled synthetic inputs, for which the ground-truth generative components are known, to natural images and videos, for which they are not. Each family exists in a *static* variant, in which an independent sample is drawn on every frame, and (where applicable) a *sequential* variant, in which the stimulus evolves continuously over time. Unless noted otherwise, inputs are presented without whitening; only the natural-image and natural-video patches are whitened (see below).

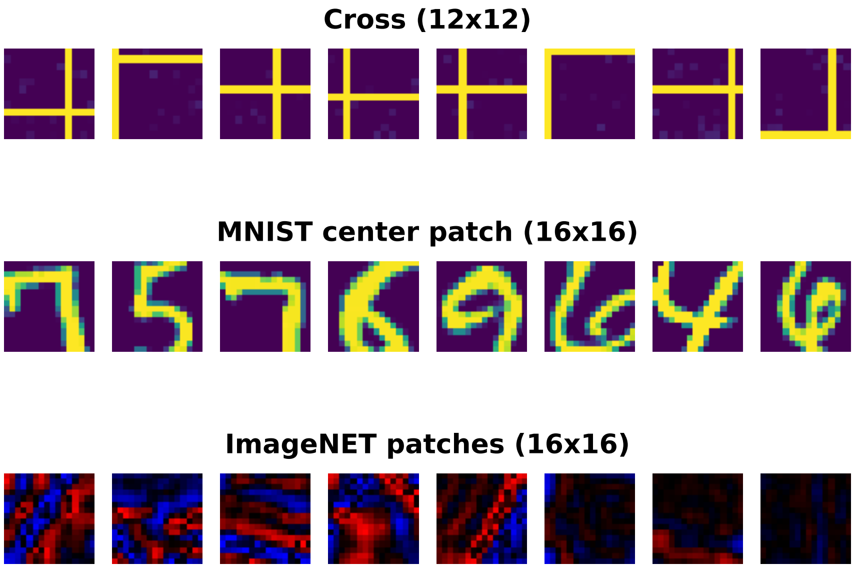


Figure S1: **Static datasets.** Examples of the static visual stimuli used to train the networks. **(Top row)** Crossed-bars stimuli of size $12 \times 12$ with sparse background noise (frequency 0.1). **(Middle row)** MNIST, central $16 \times 16$ crop with pixel intensities in $[0, 1]$. **(Bottom row)** ImageNet $16 \times 16$ gray-scale patches, ZCA-whitened and split into on/off channels (red and blue).

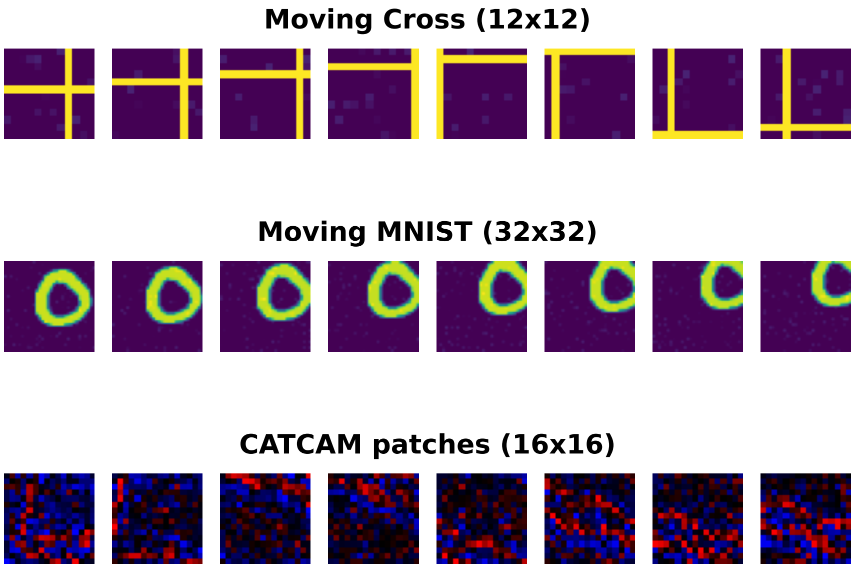


Figure S2: **Sequential datasets.** Examples of the sequential visual stimuli used to train the networks. Each row shows consecutive frames of one example sequence for each dataset. **(Top row)** Crossed-bars translating at constant velocity with wrap-around at the boundary. **(Middle row)** Moving MNIST: a $28 \times 28$ digit translating diagonally within a $32 \times 32$ frame with sparse background noise. **(Bottom row)** Natural-video patch from the CATCAM dataset: fixed $16 \times 16$ window, whitened and rectified into on/off channels.

##### Földiák bars

The bars problem [Földiak, 1990] is a standard benchmark for recovering independent generative components. Each stimulus is a $g \times g$ pixel grid on which horizontal and vertical bars are drawn at independent, uniformly random positions: each of the $2g$ possible bars is present independently. The ideal code assigns one unit per bar, so a dictionary that has recovered the generative model contains $2g$ atoms, each encoding a single bar. Because the bars activate independently, this stimulus directly exercises recovery of a known, decorrelated component set, and we use it to compare the learned dictionary against ground truth.

##### Crossed-bars

We reduce the bars problem to one horizontal and one vertical bar—a cross (Fig. S1, top). Bars are one pixel wide and drawn at unit contrast. In the *static* condition the two bars appear at independent, uniformly random positions on each frame. In the *sequential* condition each bar translates across the grid at a constant integer velocity drawn independently and uniformly from $\{-s, \ldots, -1, 1, \ldots, s\}$ pixels per frame (with $s = 1$ unless noted otherwise), wrapping around at the grid boundary to re-enter from the opposite side (Fig. S2, top). Frames are additionally corrupted by sparse background noise: each pixel is independently set, with probability $\rho = 0.1$, to a value drawn uniformly from $[0, \rho)$, and is zero otherwise (bars overwrite the noise). We vary the frame size $g \in \{12, 14, 24\}$. The generator exposes the bar positions and directions as ground-truth labels, against which the selectivity of learned units is measured.

##### Single bar

To probe orientation and direction selectivity in isolation, we further reduce the stimulus to a single bar present at any time. In this case however, the bar can have 8 different orientations, with angles evenly distributed between 0 and $\pi$. The bar is either static at a random position or translates across the grid at constant velocity in one of two directions (forward or backward) along the orientation axis. This minimal stimulus provides an unambiguous ground-truth orientation and direction for each frame, against which the tuning of learned units is measured.

##### MNIST

We use the MNIST handwritten-digit dataset [LeCun et al., 1998]. In the static condition we crop the central $16 \times 16$ region of each $28 \times 28$ image and rescale pixel intensities to $[0, 1]$ (Fig. S1, middle); the first 1,000 images are held out as a validation set. MNIST has no known ground-truth generative factors; we use it as an intermediate case of structured but naturalistic input and report reconstruction quality and the learned features.

In the sequential condition (*moving MNIST*, Fig. S2, middle) the full $28 \times 28$ digit is embedded in a $32 \times 32$ frame at an initial offset drawn uniformly from $[-5, 10)$ pixels along each axis, so the digit may initially lie partially outside the frame. The digit then translates diagonally at one pixel per frame, with the sign of the horizontal and vertical velocity components drawn independently at random for each sequence. Frames are corrupted with the same sparse background noise as the crossed-bars stimuli ($\rho = 0.1$). The generator provides the digit class and the two motion-direction components as ground-truth labels.

##### Natural images and videos

For the static condition we extract $16 \times 16$ gray-scale patches at uniformly random locations from ImageNet [Fei-Fei et al., 2009]. Patches are whitened with a zero-phase (ZCA) transform whose covariance is estimated from $10^4$ randomly sampled patches and truncated to the top 50 principal components, and the signed whitened output is split into rectified on and off channels, yielding a $16 \times 16 \times 2$ input (Fig. S1, bottom). Natural-image patches are the regime in which sparse coding is known to recover oriented, localized, bandpass features resembling V1 simple-cell receptive fields, and we use them to test whether our networks reproduce this result.

For the sequential condition we extract $16 \times 16$ patches at random spatio-temporal locations from the CATCAM video dataset [Betsch et al., 2004], spatially downsampled by a factor of 2, taking windows of 14 consecutive frames at a fixed retinal location; within a sequence, the window then advances one frame at a time, giving a temporally continuous stream (Fig. S2, bottom). Because most random natural-video patches are static, windows whose mean absolute temporal derivative falls below the 50th percentile of a random sample are rejected, so that motion is the dominant signal. Windows are whitened with a spatio-temporal ZCA transform fitted on $2 \times 10^4$ motion-selected windows (regularized in proportion to the mean eigenvalue and rescaled to unit output variance); at each time step only the most recent—causal—frame of the whitened window is read out and rectified into on/off channels, so the effective input remains a $16 \times 16 \times 2$ frame that is temporally band-passed. Sequences are split at the level of whole video clips into training, validation (8 clips), and test (8 clips) sets. As an approximate ground truth for motion, the generator estimates the local patch velocity by block matching between consecutive frames (median over frame pairs, with sub-pixel parabolic refinement), from which speed and direction labels are derived.

#### S7 Analysis

##### Reconstruction error

For each of $50$ held-out test inputs we infer the latent code $\mathbf{y}^*$, form the reconstruction $\hat{\mathbf{x}} = W^\top \mathbf{y}^*$, and report the cosine reconstruction error, defined as $1 - \cos(\mathbf{x}, \hat{\mathbf{x}})$.

We also test the model's ability to *extrapolate* a partially observed sequence. Each test sequence is presented for its first half; the second half is replaced by blank input while the network's excitatory recurrent dynamics continue to evolve the latent state. We reconstruct the full sequence from the latent trajectory and compare it to the complete ground-truth sequence, including the with held second half. Per frame we use the cosine error $1 - \cos(\mathbf{x}_t, \hat{\mathbf{x}}_t)$, averaged over frames and over $25$ test sequences of size $10$. This measures whether the learned recurrent weights have captured the temporal structure well enough to predict unseen frames, rather than merely reconstructing the current observed ones.

Finally, we test the continuous model in a similar way to the sequential model. We hold each frame for a period of time $(150\tau)$ and then take the reconstruction to be the projection of the network at the end of that time. For the empty frames, we take the projection of the network after the period of time for one frame.

##### Orientation selectivity

We characterize the orientation tuning of each unit in networks trained on ImageNet and CATCAM patches by evaluating their responses across a full range of angles. The test stimulus consists of a static bar (2 pixels in thickness) presented on a $16 \times 16$ visual field at $8$ evenly spaced orientations between $0$ and $\pi$.

Following the standard peak-response methodology in visual neuroscience [Niell and Stryker, 2008], we define the preferred orientation ($O_{\text{pref}}$) for each unit as the angle that elicits the maximum mean response. The orthogonal response ($O_{\text{ortho}}$) is calculated as the response to the orientation perpendicular ($\pm 90°$) to the preferred orientation. The orientation selectivity index (OSI) is then computed as

$$\text{OSI} = \frac{O_{\text{pref}} - O_{\text{ortho}}}{O_{\text{pref}} + O_{\text{ortho}}}. \tag{S48}$$

Assuming non-negative unit activations, this value ranges between $[0, 1]$. A value approaching $1$ indicates highly specialized tuning for a specific orientation, whereas a value near $0$ indicates equal responsiveness across orthogonal orientations. We report the OSI across units for each network, with units ordered by OSI, and compare the resulting distributions across models.

##### Direction selectivity

We also characterize the directional tuning of networks trained on the CATCAM patch dataset by assessing temporal responses to moving stimuli. A moving bar (2 pixels in thickness) is swept continuously across the $16 \times 16$ visual field in $16$ evenly spaced directions from $0$ to $2\pi$. For the continuous case in figure 5B, we only test $4$ evenly spaced orientations (up, right, down and left) as the networks are only trained on the moving crossed-bar dataset. The network's temporal state is reset prior to each sweep, and activity is recorded at the end of the sweep.

Using the standard direction contrast index [Niell and Stryker, 2008], the preferred direction ($R_{\text{pref}}$) is determined as the trajectory yielding the maximum time-averaged activity for each neuron. The null direction ($R_{\text{null}}$) is defined as the response to the motion exactly $180°$ opposite to the preferred direction. The direction selectivity index (DSI) is defined as

$$\text{DSI} = \frac{R_{\text{pref}} - R_{\text{null}}}{R_{\text{pref}} + R_{\text{null}}}. \tag{S49}$$

Like OSI, DSI values range between $[0, 1]$. Units that are highly selective to a specific direction of motion will exhibit a large DSI, while units that respond equally to a given trajectory and its reverse will have a DSI near $0$. We report the DSI across units for each network, with units ordered by DSI magnitude.

##### Tuning curves

To characterise the spatial tuning of individual units we sweep a single bar across the $P = 14$ positions of the stimulus grid and treat each position as a separate static inference problem. For each position $p$ the network is reset and the latent code is relaxed for $150\tau$ under the fixed frame $\mathbf{x}^{(p)}$, so that the recorded response $\mathbf{y}^{(p)}$ is the converged latent code. The tuning curve of unit $n$ is the resulting map

$$f_n(p) = y_n^{(p)}, \qquad p = 0, \ldots, P-1. \tag{S50}$$

Because the code is sparse, most units are unresponsive to any bar position, and their tuning curves are uninformative. We therefore rank units by their mean response $\bar{f}_n = \frac{1}{P}\sum_p f_n(p)$ and retain the 25 most active. Each retained unit has its own preferred position $p_n^* = \arg\max_p f_n(p)$; since we are interested in the *shape* of the tuning rather than in which position a given unit happens to prefer, we align the curves to a common centre by circularly shifting each one so that its peak lies at the origin, such that

$$\tilde{f}_n(\delta) = f_n((p_n^* + \delta) \bmod P)\,, \qquad \delta = -3, \ldots, 3. \tag{S51}$$

Then we plot the seven positions around the peak. The shift is circular, consistent with the periodic boundary of the stimulus sweep. Curves are coloured by activity rank.

The aligned overlay makes the common tuning width directly visible: the sharpness of the falloff on either side of the preferred position measures how localised each unit's receptive field is, and the consistency of the profiles across units indicates whether the network has learned a homogeneous set of localised filters or a heterogeneous mixture. We report the same measurement for each network and compare the resulting profiles.

##### Low-dimensional projection

Trained on a stimulus of period $T = 14$ frames, the excitatory recurrent operator approximates a cyclic shift of the latent code: one application advances the represented frame by one step, and $T$ applications return it to itself. The eigenvalues of such an operator are the $T$-th roots of unity, $\lambda_k = e^{2\pi i k/T}$, which lie on the unit circle in the complex plane. This motivates visualising the population state on a ring, with the discrete phase of the sequence embedded at the reference points

$$\mathbf{u}_t = \left(\cos\tfrac{2\pi t}{T},\ \sin\tfrac{2\pi t}{T}\right)^\top, \qquad t = 0, \ldots, T-1. \tag{S52}$$

We first estimate a preferred phase for each of the $N$ units from a single reference stimulus (a bar moving rightwards), started from a random position and run for $K$ epochs. At the end of epoch $i$ we read the latent state $\mathbf{y}^{(i)} \in \mathbb{R}^N$ and identify the dominant unit $n_i = \arg\max_n |y_n^{(i)}|$. Each unit accumulates the circular positions of the epochs at which it dominates, giving a projection matrix $R \in \mathbb{R}^{N\times 2}$ with rows

$$R_n = \sum_{i=0}^{K-1} \mathbb{1}[n_i = n]\ \mathbf{u}_{i \bmod T}. \tag{S53}$$

Units that never dominate receive a zero row and do not contribute. Because the stimulus is periodic, a unit that reliably dominates at the same phase on every cycle accumulates vectors that add constructively and acquires a long, sharply defined preferred phase, whereas a unit that is dominant at inconsistent phases receives contributions that partially cancel. The length of $P_n$ therefore encodes the reliability of the unit's phase tuning, as in the standard circular-mean construction, and downweights units without consistent temporal tuning.

Holding $R$ fixed, we present each of the four motion directions in turn and record the latent trajectory $\mathbf{y}(t)$ at the full temporal resolution of the dynamics, i.e. including the within-epoch relaxation and not only the end-of-epoch states used in Eq. (S53). Each state is mapped to the plane by the population-vector readout

$$\mathbf{z}(t) = R^\top \mathbf{y}(t) \in \mathbb{R}^2. \tag{S54}$$

We discard an initial transient and plot the remaining trajectory. Two properties of this construction are worth emphasising. First, $R$ is a fixed linear map, so the picture is a genuine linear projection of the population state and not a nonlinear embedding that could manufacture ring structure. Second, $R$ is estimated *only* from the rightward sequence; the other three directions are projected through the same map. Any structure they display therefore reflects their geometric relationship to the rightward ring — whether they traverse it, traverse it in reverse, or occupy a distinct subspace that collapses towards the origin — rather than a coordinate system fitted separately to each condition.

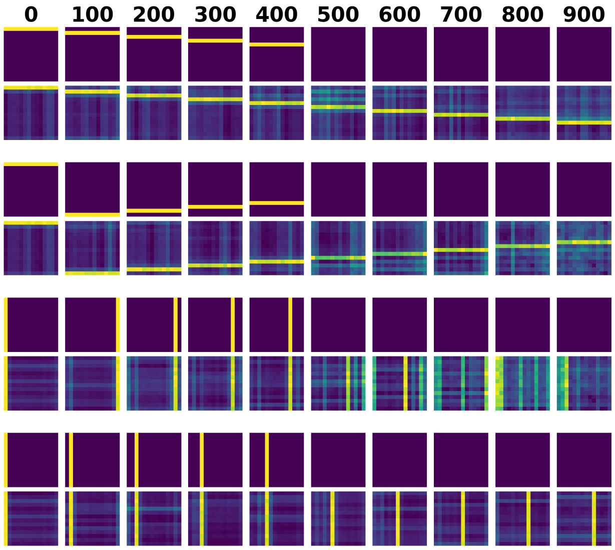


Figure S3: Examples of sequence completion in the continuous model, trained for $1,000,000\tau$ steps, using Hebbian plasticity for recurrent inhibition (matrices $M$, $U$ and $V$) and BCM plasticity for input excitation (matrices $W$ and $P$).

#### S8 Simulation details

**Network sizes.** We used 2 units for the first example with two input units (Fig. 2A), 16 units for the linear Gaussian crossbar experiment (Fig. 2B-D) and varied the number of units for different screen sizes (Fig. 2E) in order to match the input size. For the non-Gaussian case, we used 500 units across datasets. For the sequential case, we used 500 units for the moving cross and 2000 units for both moving MNIST and natural video. In the continuous case, we simulated a network with 500 excitatory neurons and 100 inhibitory neurons.

**Learning.** For the Gaussian priors, we use a fixed learning rate of $10^{-4}$, while all other cases were trained with a fixed learning rate of $10^{-3}$ across all weights. Feed-forward weights $W$ and recurrent inhibitory weights $M$ were both initialized as random matrices with entries drawn i.i.d. from $\mathcal{N}(0, 1)$. Recurrent excitatory weights were always initialized to be zero matrices. For the Gaussian case, networks were trained for 300 thousand iterations. For the non-Gaussian case, they were trained for $15,000$ iterations, and the best consistency and reconstruction scores were plotted. For the sequential case, networks were trained for 50 thousand iterations.

**Inference.** For Gaussian priors we computed the stationary latent state in closed form from Eq. S11. For non-Gaussian priors, for which no closed form exists, we obtained the stationary state by simulating the non-linear inference dynamics of Eq. S15 with step size 0.01 for 1000 steps, which is sufficient for convergence when the lateral inhibition has learned a good estimate of the second moment. If inhibition diverges, then 1000 steps still keeps the network activity within reasonable values which allows it to estimate an approximate of the second moment and eventually become stable.

**Continuous model.** To simulate the continuous model, we show moving cross sequences similar to the ones presented to the discrete model. However, in this case, each frame of the sequence is held for a period of $150\tau$. The parameters of the continuous model are set such that the recurrent excitation brings activity from the previous frame. Therefore, we set memory size $\beta = 200\tau$, the kernel mean to be $\mu = 0.5$ and the deviation to be $\sigma = 0.2$. After training for $25,000,000\tau$ we observe sequence completion abilities emerging (Fig. S3).

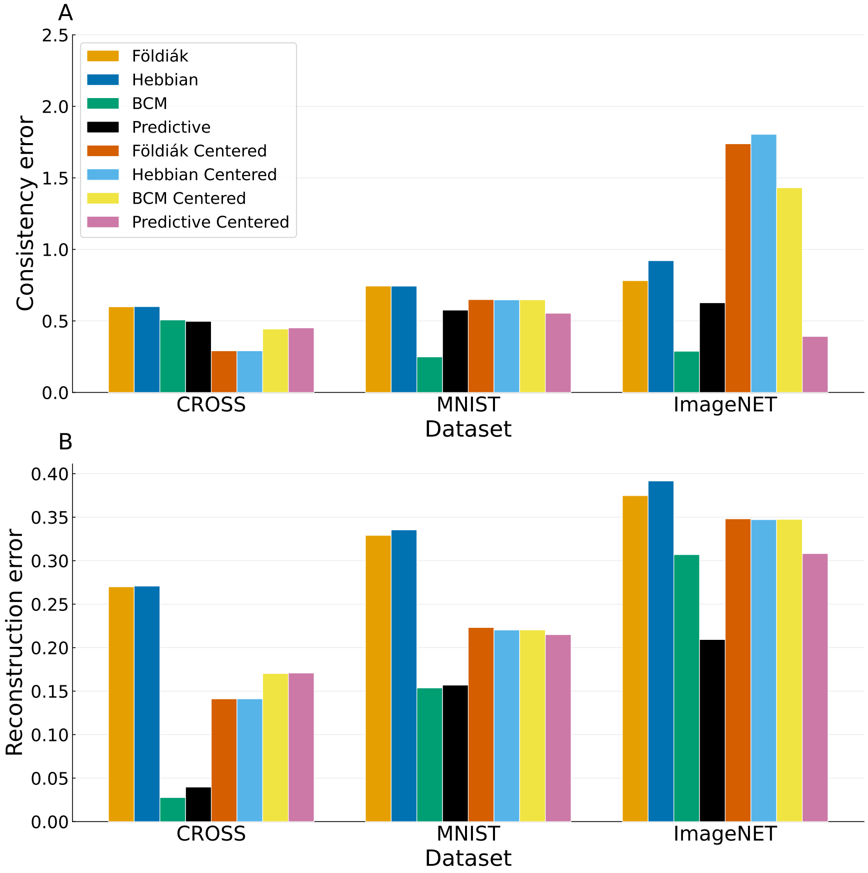


Figure S4: **Consistency and reconstruction errors for different feed-forward rules** (A) Normalized consistency error (Eqn. S4) for different datasets and feed-forward plasticity rules. Experiments were run with 5 different seeds, the mean is plotted and the variability is on average $4.9 \times 10^{-7}$ (omitted in plots). (B) Reconstruction error between the input $\mathbf{x}$ and the reconstruction from the stable state $\hat{\mathbf{x}} = W^\top \mathbf{y}^*$. Experiments were run with 5 different seeds, the mean is plotted and the variability is on average $4.7 \times 10^{-5}$ across rules.

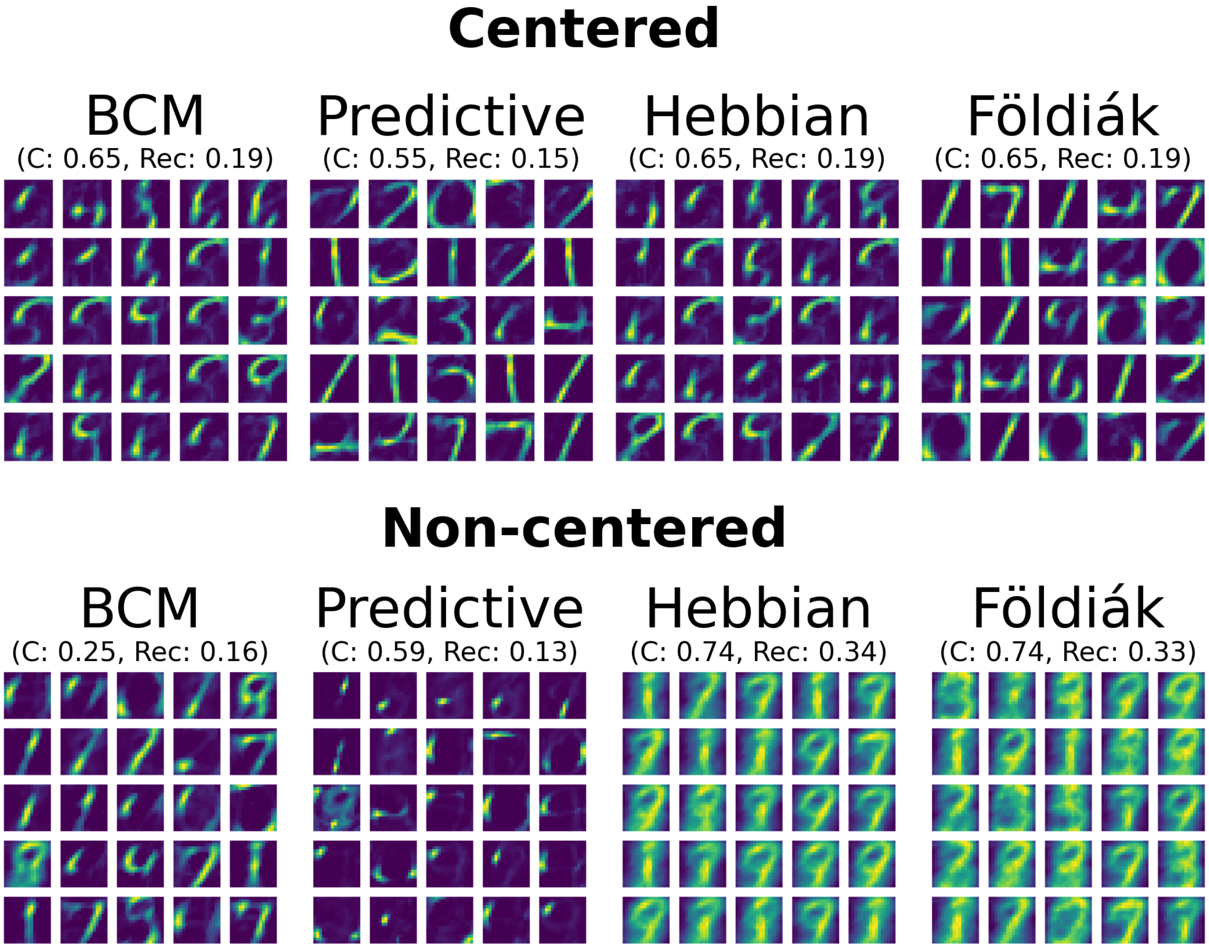


Figure S5: **MNIST receptive fields** Examples of receptive fields developed under the MNIST dataset for different feed-forward rules. All networks follow non-linear dynamics, use Hebbian plasticity for inhibition, have $500$ units, receive input normalized and keep feed-forward weights normalized. For the top row, we plot networks whose input has been centered such that $\mu_{\mathbf{x}} = \mathbf{0}$. In the bottom row we plot networks whose input is strictly non-negative, with each variable having larger than zero mean. Receptive fields, reconstruction error (Rec) and normalized consistency error (C) are plotted for networks developed under different feed-forward rules.

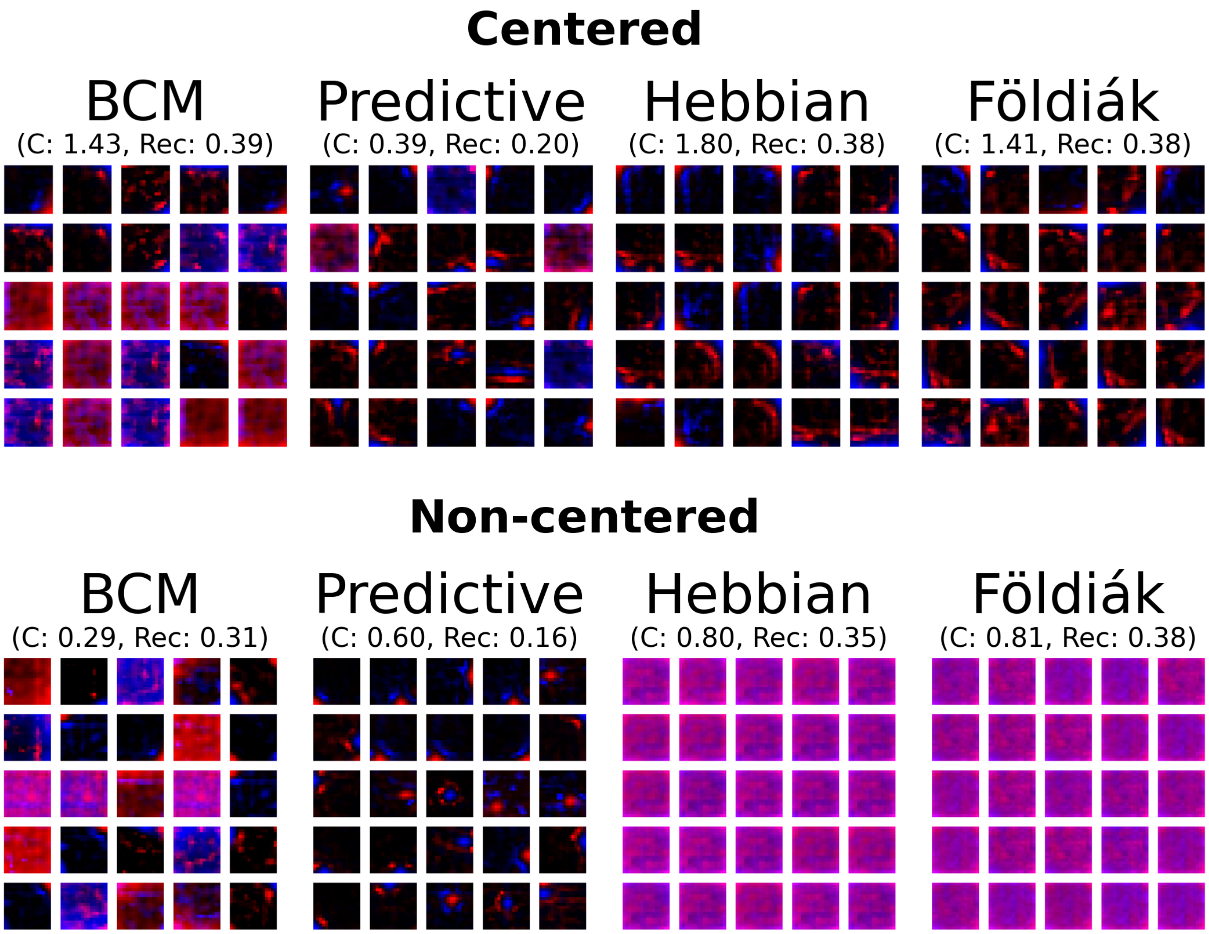


Figure S6: **ImageNET patches receptive fields** Examples of receptive fields developed under the ImageNET dataset for different feed-forward rules. All networks follow non-linear dynamics, use Hebbian plasticity for inhibition, have 500 units, receive input normalized and keep feed-forward weights normalized. For the top row, we plot networks whose input has been centered such that $\mu_{\mathbf{x}} = \mathbf{0}$. In the bottom row we plot networks whose input is strictly non-negative, with each variable having larger than zero mean. Receptive fields, reconstruction error (Rec) and normalized consistency error (C) are plotted for networks developed under different feed-forward rules.
